# ATG5 selectively engages virus-tethered BST2/Tetherin in an LC3C-associated pathway

**DOI:** 10.1101/2023.01.06.522978

**Authors:** Delphine Judith, Margaux Versapuech, Fabienne Bejjani, Marjory Palaric, Pauline Verlhac, Aurelia Kuster, Leslie Lepont, Sarah Gallois-Montbrun, Katy Janvier, Clarisse Berlioz-Torrent

## Abstract

BST2/Tetherin is a restriction factor that reduces HIV-1 dissemination by tethering virus at the cell surface. BST2 also acts as a sensor of HIV-1 budding, establishing a cellular anti-viral state. The HIV-1 Vpu protein antagonizes BST2 antiviral functions, notably by subverting an LC3C-associated pathway, a key cell intrinsic anti-microbial mechanism. Here, we show that ATG5 associates with BST2 and acts as a signaling scaffold to trigger an LC3C-associated pathway induced by HIV-1 infection. This process is initiated at the plasma membrane through the recognition of virus-tethered BST2 by ATG5. ATG5 and BST2 assemble as a complex, independently of the viral protein Vpu and ahead of the recruitment of the ATG protein LC3C. The conjugation of ATG5 with ATG12 is dispensable for this interaction. ATG5 recognizes cysteine-linked homodimerized BST2 and specifically engages phosphorylated BST2 tethering viruses at the plasma membrane, in an LC3C-associated pathway. We also found that this LC3C-associated pathway is used by Vpu to attenuate the inflammatory responses mediated by virion retention. Overall, we highlight that by targeting BST2 tethering viruses, ATG5 acts as a transducer of the LC3C-associated pathway induced by HIV-1 infection.

**Significance statement:** The outcome of viral infection in cells is dependent on the balance between host restriction factors and viral countermeasures. BST2/Tetherin is a restriction factor that reduces HIV-1 dissemination by tethering virions at the cell surface. Its action is counteracted by the viral protein Vpu through multiple mechanisms. Here, we describe the initial step of a non-canonical autophagic pathway, called LC3C-associated pathway, subverted by Vpu to counteract BST2 antiviral activities. We found that the autophagic protein ATG5 acts as a transducer by targeting phosphorylated and dimerized virus-tethered BST2 from cell surface to the degradation. Our discovery opens new avenue in the discovery of unconventional functions of ATG5, as an adaptor for receptor at the plasma membrane initiating an unconventional autophagy process.

## Introduction

Host restriction factors, a set of proteins with direct or indirect antiviral activity, are an important facet of the innate immune response controlling infection in a cell-intrinsic manner (1). Among them, the restriction factor BST2 (bone marrow stromal antigen 2)/Tetherin interferes with the release of diverse mammalian enveloped viruses including the human immunodeficiency virus type 1 (HIV-1) by physically trapping *de novo* formed viral particles at the surface of infected cells (2, 3). Tethered virions can be endocytosed and then targeted for lysosomal degradation (4). The antiviral activity of BST2 relies on its presence at the viral budding site and on its ability to be incorporated into budding virions, bridging virions and cellular membranes (5). Prior to its identification as an antiviral factor, BST2 was identified as an inducer of proinflammatory genes expression via NFĸB activation (6). Coupled to its viral tether activity, BST2 acts as a sensor of HIV-1 budding, establishing a cellular anti-viral state, and thus making infected cells more visible to innate and adaptative immunity (7, 8).

Viruses have thus adopted diverse strategies to antagonize this restriction, and further optimize viral release and attenuate the cellular anti-viral response. They notably encode counteracting viral proteins, such as the HIV-1 accessory protein Vpu. Vpu countermeasures against BST2 depends on a direct interaction between Vpu transmembrane domain and that of BST2 (5, 9, 10). Vpu downregulates BST2 cell surface expression (2, 3), by modifying BST2 intracellular trafficking (11-15) and accelerating ESCRT-mediated sorting of BST2 for degradation (16-19). Vpu also displaces BST2 from viral budding sites. This activity is crucial to overcome BST2 restriction on HIV-1 release. It involves hijacking of clathrin adaptor protein complexes (AP)-dependent endocytic pathways (11-13) and is independent of Vpu effect on BST2 cell surface expression (2, 3, 9, 13, 20). Moreover, Vpu attenuates the NFkB-dependent pro-inflammatory response mediated by viral-induced BST2 signaling (7, 21). Interestingly, we described a non-canonical autophagic pathway contributing to HIV-1 Vpu antagonism of the BST2 innate barrier. We reported that Vpu acts with LC3C, an autophagy protein (ATG) to favor the removal of BST2 molecules present at HIV-1 budding sites and their targeting in a single-membrane compartment for degradation. Two ATG proteins, ATG5 and Beclin 1/ATG6, but not all the components of the autophagy pathway, have been also implicated in this process, supporting the view that a non-canonical autophagy process reminiscent of LC3-associated pathway contributes to Vpu counteraction of BST2 restriction.

The LC3-associated pathway is a non-canonical autophagy process controlled by a subset of ATG proteins, linking activation of a surface receptor with phagocytosis (LAP) (22) or endocytosis (LANDO) (23). LC3-associated pathway has a prominent anti-inflammatory function and represents a key cell intrinsic restriction mechanism against infections (24-26). As expected for such an anti-microbial pathway, pathogens have acquired the ability to subvert it to attenuate the host responses or establish a proliferative niche in the host (26, 27). LC3-associated pathway differs from canonical autophagy by the formation of a single-membrane structure from invagination of the plasma membrane (28). The engagement of the surface receptor triggers the recruitment of several components of the canonical autophagy process to conjugate LC3 at the phagosomal or endosomal membrane (25, 29). This recruitment accelerates the fusion of these single membrane compartments with the lysosomes and thus the degradation of their contents. In this process, ATG5 in conjugation with ATG12 is part of the LC3 conjugation system which catalyzes the conjugation of LC3 on the single membrane vesicle. Although the molecular players involved in the initiation and regulation of LC3-associated phagocytosis are becoming better characterized, the molecular mechanism driving the selective engagement of BST2 molecules in the LC3-associated pathway induced by HIV-1 infection are not elucidated (29-31). In this study, we dissected the initial events driving BST2 in the LC3C-associated pathway subverted by Vpu and analyzed the contribution of this pathway on Vpu countermeasure to pro-inflammatory signaling induced by BST2 sensing.

Here, we identify ATG5 as a key player in the LC3C-associated pathway subverted by Vpu. We show that the formation of an ATG5 and BST2 complex at the cell surface is the first step of this viral-induced LC3C-associated process. It occurs independently of Vpu, ahead of the recruitment of LC3C, and requires cysteine-linked homodimerization of BST2. We revealed that the unconjugated form of ATG5 can efficiently recruit the dimerized form of BST2. We also provide evidence that ATG5 preferentially targets phosphorylated virus-tethered BST2, the specific form triggering BST2 signaling. Finally, we found that this LC3C-associated pathway is used by Vpu to attenuate the inflammatory responses mediated by BST2 sensing of virion retention. In summary, we propose that by targeting BST2 tethering viruses, ATG5 acts as a transducer of the LC3C-associated pathway induced by HIV-1 infection.

## Results

### ATG5 and BST2 assemble as a complex independently of Vpu and ahead of the recruitment of LC3C

We previously described a non-canonical autophagy pathway reminiscent of LC3-associated phagocytosis contributing to HIV-1 Vpu antagonism of the BST2 innate barrier. We highlighted that ATG5 and Beclin 1, but not all the components of the autophagic pathway, act with LC3C in this process. We revealed a selective interaction between HIV-1 Vpu and LC3C (Figure 1A)(27).

**Figure 1.**
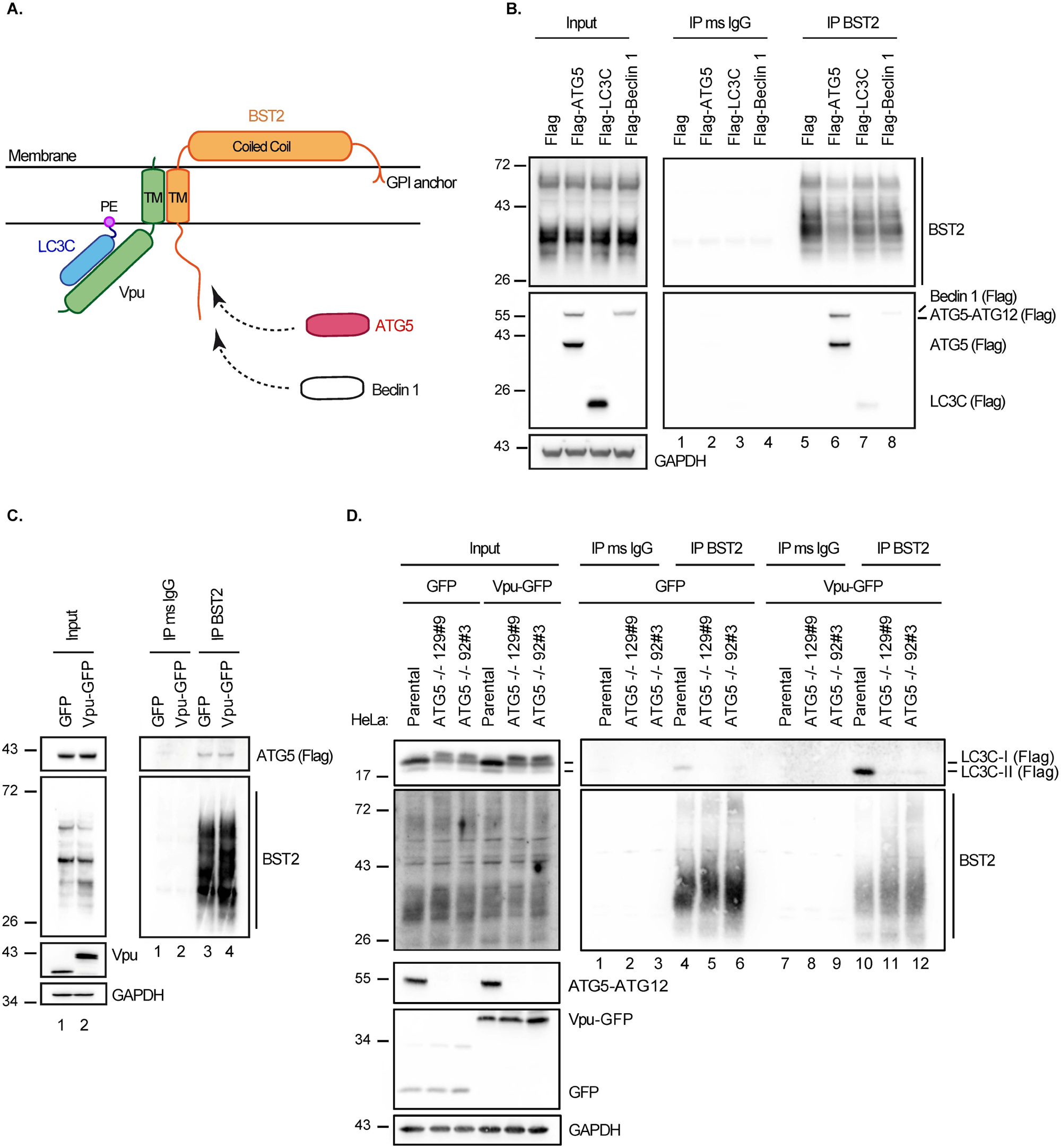
ATG5-BST2 complex forms independently of Vpu and ahead of LC3C recruitment. **(A)** Overview of the interaction between BST2, Vpu, LC3C and ATG5. **(B)** Immunoprecipitation of endogenous BST2 in HeLa cells extracts co-transfected with plasmids expressing Flag-ATG5, Flag-LC3C and Flag-Beclin 1. Immunoprecipitated proteins were detected by western blotting using rabbit anti-BST2 and anti-Flag HRP antibodies. **(C)** Immunoprecipitation of endogenous BST2 in HeLa cells extracts co-transfected with plasmids expressing Flag-ATG5 and GFP or Vpu-GFP. Immunoprecipitated proteins were detected by western blotting using rabbit anti-BST2 and anti-Flag HRP antibodies. **(D)** Immunoprecipitation of endogenous BST2 in Parental (CTRL) or ATG5 knockout (ATG5 -/- 129#9 or 92#3) HeLa cells extracts co-transfected with vectors encoding for Flag-LC3C and GFP or Vpu-GFP. Immunoprecipitated proteins were detected by western blotting using rabbit anti-BST2 and anti-Flag-HRP antibodies. All Western blots presented are representative of at least three independent experiments. Immunoprecipitation of endogenous BST2 were done with mouse anti-BST2 antibodies in all experiments.

To determine the initial step driving the engagement of BST2 molecules present at the budding site in the LC3C-associated pathway induced by HIV-1 infection, we examined whether BST2 interacts with ATG5, Beclin 1 or LC3C. We found through co-immunoprecipitation experiments that BST2 interacts with Beclin 1 and LC3C albeit with low affinity (figure 1B, compare lanes 7-8 to lane 5). By contrast, we revealed a strong interaction between BST2 and the two forms of ATG5, the unconjugated form of ATG5 (ATG5) and the ATG5 form conjugated to ATG12 (ATG5-ATG12) (figure 1B, compare lanes 5 to 6). Moreover, we observed that Vpu expression did not modify this interaction (Figure 1C, right panel, compare lanes 3 to 4), suggesting that ATG5 interacts with BST2 independently of Vpu and of its conjugation to ATG12.

We previously showed that, upon infection, Vpu binds specifically to the ATG protein LC3C. This interaction favors the targeting of BST2 molecules present at the budding site to an LC3C-associated pathway (27). The complex made by LC3C, Vpu and BST2 plays a central part in this process (Figure 1A). We thus analyzed the contribution of LC3C in the interaction between BST2 and ATG5. Using two LC3C knockout cell lines previously described in (27), we observed that the presence of LC3C is not required for ATG5 to bind BST2 (Supplemental figure 1A, right panel, compare lanes 5-6 to 4). Conversely, we examined whether the formation of the complex made by LC3C, Vpu and BST2 relies on the expression of ATG5. For that, we established two different ATG5-/- HeLa clones (ATG5-/- 129#9 and 92#3) using the CRISPR-Cas9 system. As expected, the knock-out (KO) of ATG5 gene blocked the lipidation of LC3 proteins and only the non lipidated form of LC3C (LC3C-I) was observed even upon autophagy induction by amino-acid starvation (ES) (Supplemental figure 1B). Using co-immunoprecipitation, we showed that only the lipidated form of LC3C (LC3C-II) binds weakly to BST2 in presence of ATG5 and this binding is strongly increased in presence of Vpu, correlating with a Vpu-dependent co-immunoprecipitation of LC3C with BST2 (Figure 1D, right panel, compare lanes 4 to 10) (27). In contrast, the knock-out of ATG5 resulted in a loss of binding of LC3C to BST2 even upon Vpu expression (Figure 1D, right panel, compare lanes 11-12 to 10), indicating that ATG5 is a key determinant for the formation of this 3-partner complex.

Altogether, these data support the view that ATG5 and BST2 assemble as a complex independently of Vpu and ahead of the recruitment of LC3C.

### ATG5 takes in charge BST2 molecules tethering viral particles at the cell surface

To confirm the implication of ATG5 in the engagement of BST2 molecules in the LC3C-associated pathway induced by HIV-1 infection, we assessed the impact of ATG5 depletion on the distribution of BST2 and viral components at the surface of HIV-1 infected cells. Confocal analysis of cell surface BST2 staining revealed rare small BST2 puncta at the surface of control infected cells (Figure 2A, upper panel) compared to BST2 staining in non-infected cells, illustrating the well documented BST2 down-regulation mediated by Vpu (2). Interestingly, in ATG5-depleted cells, large puncta of BST2 were still observed in specific area of the plasma membrane, colocalizing with HIV structural components, the tagged HIV-1 Gag product MA/YFP and the HIV-1 Envelope glycoprotein (Env) (Figure 2A), despite an overall down regulation of cell surface BST2 in infected cells compared to neighboring non-infected cells. Indeed, measurement of BST2 cell surface level by flow cytometry showed that Vpu-mediated down-regulation of cell surface BST2 is still efficient in ATG5-depleted cells infected with HIV-1 WT (CAp24 positive cells) compared to non-infected cells or cells infected with HIV-1 that do not express Vpu (HIV-1 Udel) (Supplemental figure 2).

**Figure 2.**
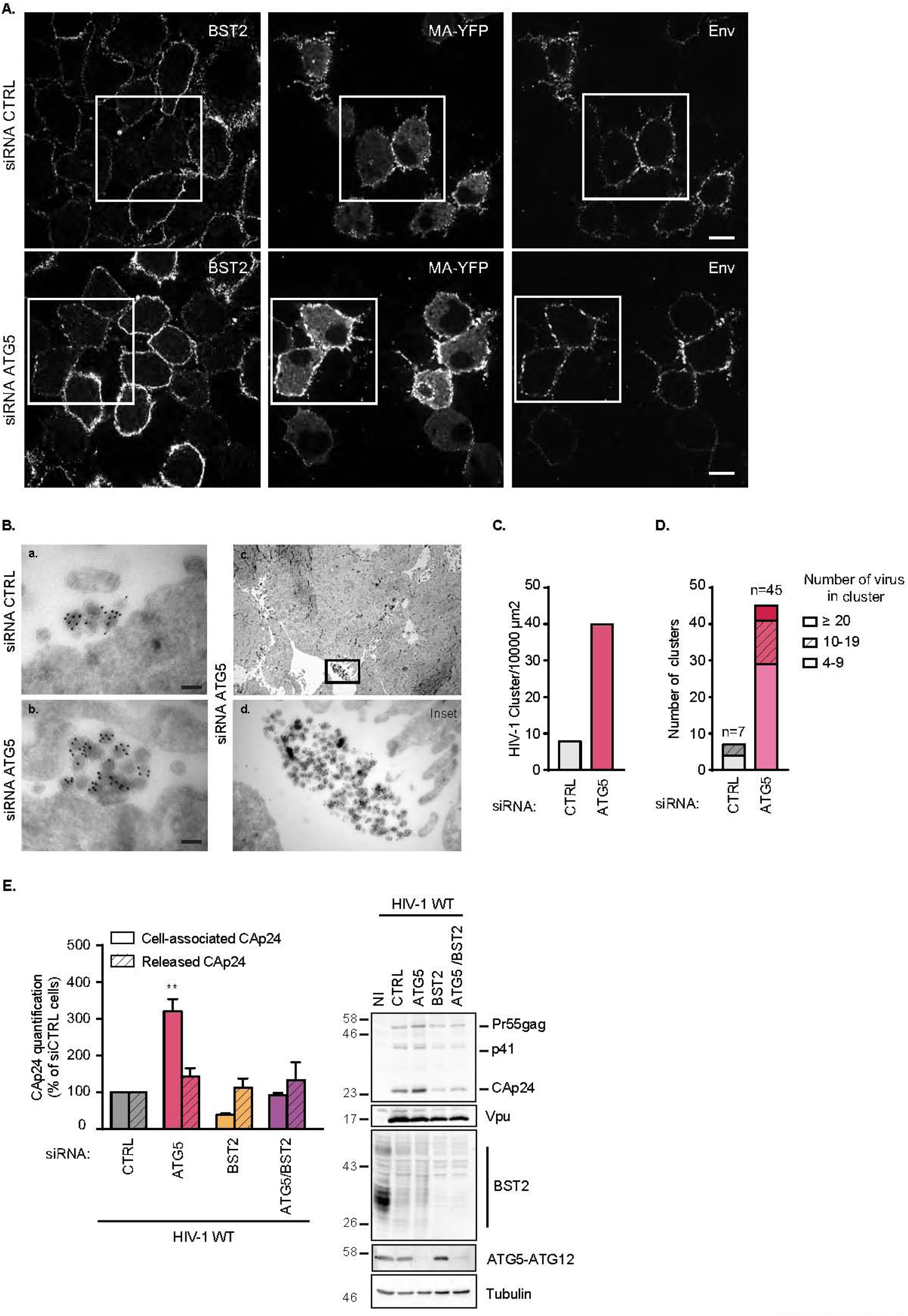
ATG5 depletion leads to virions retention at the plasma membrane in a BST2 dependent manner. **(A)** Confocal fluorescence microscopy of HeLa cells transfected with indicated siRNA and infected with VSV-G-pseudotyped HIV-1 WT MA-YFP at a MOI of 0.5 for 24 hrs following cell surface staining with human anti-Env (SUgp120) and rabbit anti-BST2 antibodies. Scale Bar: 10µM. Immunofluorescence images presented are representative of at least three independent experiments. **(B)** Cryosections of HeLa cells treated with CTRL (a) or ATG5 (b) siRNA for 72hrs and then infected with VSV-G pseudotyped HIV-1 WT for 48 hrs before labelling with antibodies against CAp24 and 10-nm protein A-gold. Scale bar, 100nm. (c) Clusters of viral particles at the cell surface in ATG5 depleted cells. (d) Enlargement of the region indicated in the upper panel (c). **(C)** Quantification of the number of clusters of viral particles per 10000 µm^2^ analyzed in (A). The number of clusters of viral particles in control, ATG5 and Beclin-1-depleted cells was determined by scanning 8815, 11540 and 10315 µm^2^, respectively. **(D)** Relative distribution of HIV-1 particles in cluster presented in (A). The total number of clusters of HIV-1 particles counted in control, ATG5 and Beclin-1-depleted cells was 7, 45 and 15, respectively. **(E)** HeLa cells transfected with indicated siRNA were infected with a VSV-G pseudotyped HIV-1 NL4.3 WT at a MOI of 0.5 for 48 hrs. Cell-associated CAp24 and released CAp24 were quantified by Elisa. Statistical analysis using two-way ANOVA with Holm-Sidak’s multiple comparison test, mean ± SEM, n=3 experiments; *p ≤ 0.05, ** p ≤ 0.01. Western blot analysis of HIV-1 Gag and CAp24 products, Vpu, BST2, ATG5 and tubulin in infected siRNA-treated cells. All western blot are representative of at least three independent experiments.

We then investigated the nature of the large patches of viral products accumulated at the periphery of ATG5-depleted cells by electron microscopy. Labeling of cryosections of HIV-1 WT infected cells with antibody targeting the viral Gag subproduct CAp24 (Capsid) mainly showed mature HIV-1 particles accumulating at the plasma membrane in ATG5-depleted cells (Figure 2B, panel b). Analysis of HIV-1 particles distribution revealed a higher number of HIV-1 particles clusters per 10000 µm^2^ found at the cell surface in ATG5-depleted cells compared to infected control cells (Figure 2B, panels c-d and Figure 2C). Furthermore, the average number of HIV-1 particles per cluster upon ATG5 depletion was superior to that in control cells (Figure 2D). Interestingly, while we saw numerous intracellular structures containing mature HIV-1 particles in LC3C-depleted cells in our previous study (27), we rarely observed HIV-1 particles sequestered in intracellular compartments in ATG5 depleted cells. These data provide evidence that, upon HIV-1 infection, ATG5 acts at the cell surface level and upstream of LC3C.

We finally checked whether the effect of ATG5 depletion on HIV-1 particle retention at the cell surface was related to BST2 expression. Upon HIV-1 infection, ATG5 depletion induced a strong accumulation of the cell-associated viral CAp24 protein and did not impact the amount of CAp24 released found in the supernatant. This effect can also be detected by western blot in ATG5-depleted infected cells (figure 2E). Interestingly, ATG5 knockdown did not induce an accumulation of cell-associated CAp24 nor affected the amount of CAp24 released in BST2-depleted cells suggesting, along with the cell microscopy data, that the effect of ATG5 silencing on viral retention is dependent of BST2.

Together, these results suggest that ATG5 takes in charge BST2 molecules tethering viral particles at the cell surface.

### The interaction between ATG5 and BST2 is strengthened in HIV-1 producing cells and independent of ATG5 conjugation status

Based on the above observations (Figure 2), we hypothesized that ATG5 acts as an adaptor to trigger the targeting of virus-tethered BST2 onto the LC3C-associated pathway subverted by Vpu. We thus explored the binding of ATG5 on BST2 in HIV-1 producing cells. We found through co-immunoprecipitation that the interaction between ATG5 and BST2 is strengthened upon HIV-1 production (Figure 3A). Interestingly, we noticed that HIV-1 productive infection favors the recruitment of unconjugated form of ATG5 on BST2, independently of Vpu expression (Figure 3A, right panel, compare lanes 4 and 6 to 2). This correlates with the Vpu-independent binding of ATG5 to BST2 shown in Figure 1C.

**Figure 3.**
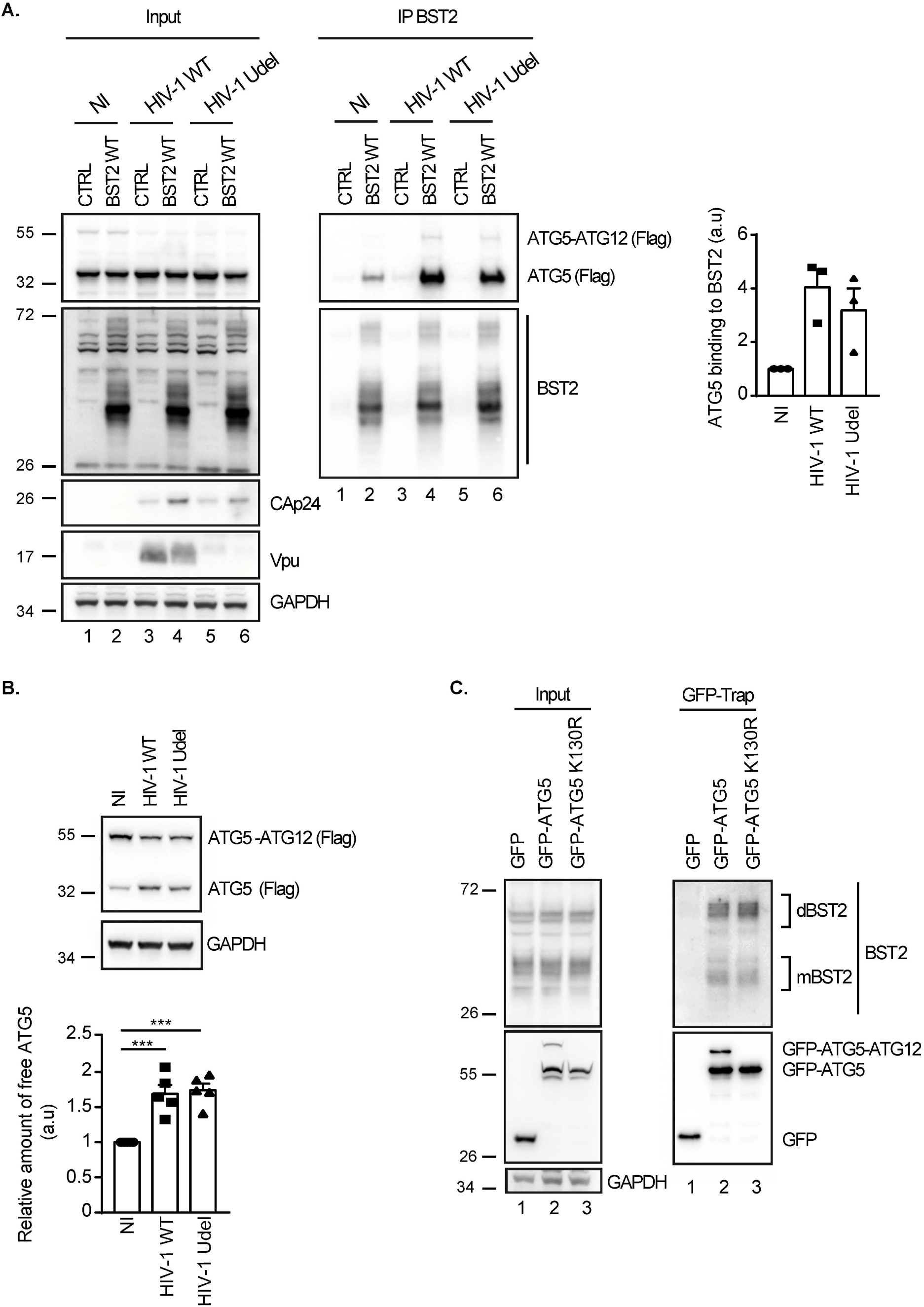
Productive infection favors ATG5-BST2 interaction independently on ATG5 conjugation with ATG12. **(A)** HEK293T cells were co-transfected with p3XFlag-ATG5, either pcDNA or pcDNA-BST2 WT and the provirus HIV-1 WT or Vpu-deleted (Udel) for 24 hrs. BST2 was immunoprecipitated with mouse anti-BST2 antibodies and immunoprecipitated proteins were detected by western blotting using rabbit anti-BST2 and anti-Flag-HRP antibodies. Quantification of the relative amount of ATG5 immunoprecipitated with BST2 WT upon infection in three experiments. **(B)** HeLa cells were co-transfected with vector encoding for Flag-ATG5 and the provirus HIV-1 WT or Vpu-deleted (Udel) for 24 hrs. Proteins were detected by western blotting using anti-Flag-HRP and anti-GAPDH antibodies. Quantification of the relative amount of unconjugated ATG5 after normalization to the ATG5-ATG12 conjugate form upon infection in five experiments. **(C)** Immunoprecipitation of GFP in HeLa cells extracts transfected with vectors expressing GFP, GFP-ATG5 or GFP-ATG5 K130R. Immunoprecipitated proteins were detected by western blotting using rabbit anti-BST2 and anti-GFP HRP antibodies.

As unconjugated ATG5 binds BST2, we next studied the conjugation pattern of ATG5 upon infection. Intriguingly, we observed that independently of Vpu expression, the amount of ATG5-ATG12 conjugate ismarkedly reduced in infected cells, whereas the amount of free ATG5 increased significantly suggesting that the infection reduces the formation of ATG5-ATG12 conjugate, making free ATG5 available (Figure 3B).

Finally, we evaluated whether the binding of ATG5 to BST2 relies on its conjugation status with ATG12, and thus on its function in autophagy. We performed a green fluorescent protein affinity immunoprecipitation (GFP-Trap) in cells expressing GFP-ATG5 or GFP-ATG5 K130R, an autophagy-incompetent mutant bearing a point mutation that prevents the formation of the ATG5-ATG12 conjugate essential in autophagy. The results showed that GFP-ATG5 WT or GFP-ATG5 K130R similarly immunoprecipitates BST2. Interestingly, both constructs bound predominantly the dimeric form of BST2 (Figure 3C).

Thus, these data suggest that ATG5-ATG12 conjugation is dispensable for the interaction between ATG5 and BST2 and that ATG5 could bind preferentially dimeric forms of BST2.

### Binding of ATG5 to the cytosolic domain of BST2 requires BST2 dimerization

To further understand the initial event that governs the recruitment of ATG5 on BST2, we investigated which domain of BST2 interacts with ATG5 and spotted that ATG5 binds to the cytoplasmic domain of BST2, since the truncation of the first 21 amino-acids of BST2 strongly decreases ATG5-BST2 interaction (Figure 4A, compare lanes 2 to 3). To confirm our finding, we performed an immunoprecipitation assay using the short isoform of BST2 lacking the first 12 amino acids of the cytoplasmic tail (BST2 M1A) (32) (Supplemental figure 3A). We observed that the interaction between ATG5 and the short isoform of BST2 (BST2 M1A) is less efficient than with BST2 WT, indicating that the integrity of the cytoplasmic tail of BST2 is necessary for its interaction with ATG5. We next performed an alanine scanning mutagenesis along the cytoplasmic domain of BST2 to determine the minimal sequence required for the binding of ATG5 to BST2 (Supplemental figures 3B and 3C). Interestingly, none of the mutants affected the binding of ATG5 to BST2, implying that current mutations are not enough to alter the motif or structure essential for the ATG5-BST2 interaction.

**Figure 4.**
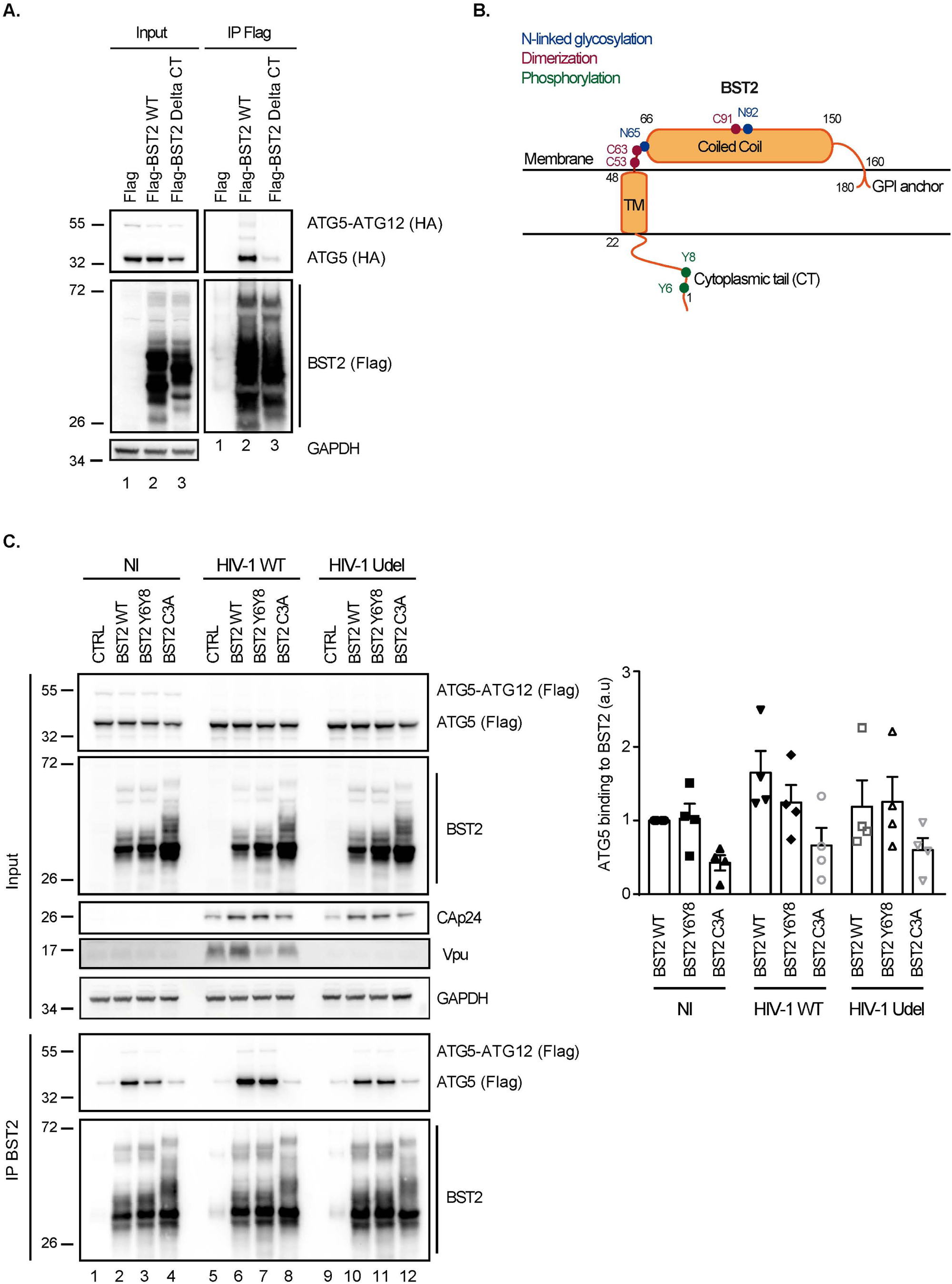
BST2 dimerization is an essential for efficient ATG5 binding. **(A)** HeLa cells were transfected with plasmids expressing HA-ATG5 and either Flag, Flag-BST2 or Flag-BST2 truncated for its cytosolic domain (BST2 Delta CT). Flag-tagged proteins were immunoprecipitated with anti-Flag antibodies. Immunoprecipitated proteins were detected by western blotting using anti-HA-HRP and anti-Flag-HRP antibodies. **(B)** Schematic representation of human BST2 showing the GPI anchor, the coiled-coil and the transmembrane domains as well as the cytoplasmic tail. Indicated in blue, red and green respectively were the amino acids essential for the N-linked glycosylation (N_65_ and N_92_), dimerization (C_53_, C_63_, C_91_) and phosphorylation (Y_6_, Y_8_) of BST2. **(C)** HEK293T cells were co-transfected with vectors expressing Flag-ATG5, either pcDNA-BST2 WT or BST2 Y_6_Y_8_ (double tyrosine phosphorylation defective mutant) or BST2 C3A (triple cysteine dimerization defective mutant) and the provirus WT or Vpu-deleted (Udel) NL4.3 HIV-1 for 24 hrs. BST2 was immunoprecipitated with mouse anti-BST2 antibodies and immunoprecipitated proteins were detected by western blotting using rabbit anti-BST2 and anti-Flag-HRP antibodies. Quantification of the relative amount of ATG5 immunoprecipitated with BST2 WT; BST2 Y_6_Y_8_ or BST2 C3A upon infection in four experiments. All Western blots presented are representative of at least three independent experiments.

Upon infection, BST2 acts as a viral tether *via* multimerization by bridging viral particles to the cells as well as a viral sensor capable of initiating the recruitment of proteins and signaling cascades upon infection (7, 21). We thus investigated whether mutations that disrupt the sensing and tethering function of BST2 affect the binding of ATG5. To alter the sensing function of BST2, we mutated the 2 conserved tyrosine residues (Y_6_ and Y_8_) that are phosphorylated upon retroviral particle retention (8). This BST2 Y_6_Y_8_-AA mutant (BST2 Y_6_Y_8_) is unable to activate the downstream NFĸB pathway (Figure 4B) (7, 21, 32). Accordingly, BST2 Y_6_Y_8_ was not recognized by an antibody that we developed, targeting specifically phosphorylated forms of BST2 (pBST2) (Supplemental figure 3D, compare lanes 2 to 3). For the tethering function of BST2, we focused on BST2 C3A mutant in which three cysteines (C_53_, C_63_ and C_91_) of BST2 ectodomain required for dimerization have been mutated to alanine (Figure 4B). This mutant is unable to dimerize and to tether virions at the cell surface (33). Migration of this mutant under non-reducing condition confirmed its inability to dimerize (Supplemental figure 3E, compare lanes 2 to 4). Co-immunoprecipitation experiments showed that the substitution of Y_6_Y_8_ residues to alanine residues (BST2 Y_6_Y_8_) did not alter ATG5-BST2 interaction, indicating that phosphorylation of the cytoplasmic tail is not required for ATG5 interaction (Figure 4C, compare lanes 2 to 3, lanes 6 to 7 and lanes 10 to 11). Interestingly, we found that the mutation of the 3 cysteines of BST2 ectodomain (BST2 C3A) decreases the binding of ATG5 to BST2 (Figure 4C, right panel, compare lanes 2 to 4, lanes 6 to 8 and lanes 10 to 12).

Altogether, these results indicate that ATG5 binding to BST2 requires the integrity of the cytoplasmic tail of BST2 as well as the cysteine residues implicated in BST2 dimerization and virus tethering.

### ATG5 targets phosphorylated dimerized virus-tethered BST2

To further define the form of BST2 targeted by ATG5 upon tethering of viral particles, we determined the impact of ATG5 depletion on the N-linked glycosylation pattern of BST2 upon infection. BST2 is a N-linked glycosylated protein on asparagine 65 and 92 (N_65_; N_92_) (Figure 4B) that can be found in a monomeric (mBST2) and dimeric form (dBST2). Following crosslinking with formaldehyde, allowing us to study simultaneously monomeric and dimeric status of BST2, mBST2 migrates as a smear of heterogeneously glycosylated forms with apparent molecular weights between 26-43kDa in control cells, whereas dBST2 migrates between 43-95 kDa (Supplemental figure 4A, lane 1). Several glycosylated forms of BST2 can be detected: the non-glycosylated species, the high mannose modification, and complex-type carbohydrates at either or both positions. Interestingly, ATG5 depletion induced an upward shift of BST2 smear for both mBST2 and dBST2, compared to control cells or LC3C depleted cells (Supplemental figure 4A, lanes 2, 5 and 8 to lanes 1, 3 and 4, and to lanes 6, 7 and 9), suggesting that the N-linked glycosylated forms of BST2 are enriched in absence of ATG5. As similar results were observed in cells infected with HIV-1 WT and Udel compared to non-infected cells, this phenotype was independent of the infection and the expression of Vpu (Supplemental figure 4A, compare lanes 5 and 8 to lane 2; see also Supplemental figure 4B, line scan HIV-1 WT vs HIV-1 Udel).

To go further, we assessed the impact of ATG5 depletion on the dimerization of BST2. A PNGase treatment, which removes all oligosaccharides on BST2 molecules enriched after immunoprecipitation, resulted in deglycosylated mBST2 with a molecular weight of 17kDa and a deglycosylated dBST2 with a molecular weight of 38kDa (Supplemental figure 4C, compare Lane 1 to lane 4). By evaluating the ratio between dBST2 and mBST2 to assess the variations of BST2 expression in each condition, we revealed an increased amount of dBST2 in ATG5 depleted cells compared to control cells, in both infected and non-infected cells (Figure 5A, upper panel, compare lanes 2,5 and 8 to lanes 1, 4 and 7 and Figure 5B). These results suggest that, upon infection, ATG5 targets virus-tethered BST2 which is mainly N-linked glycosylated and under its dimeric form, confirming the results shown in the figures 3C and 4C.

**Figure 5.**
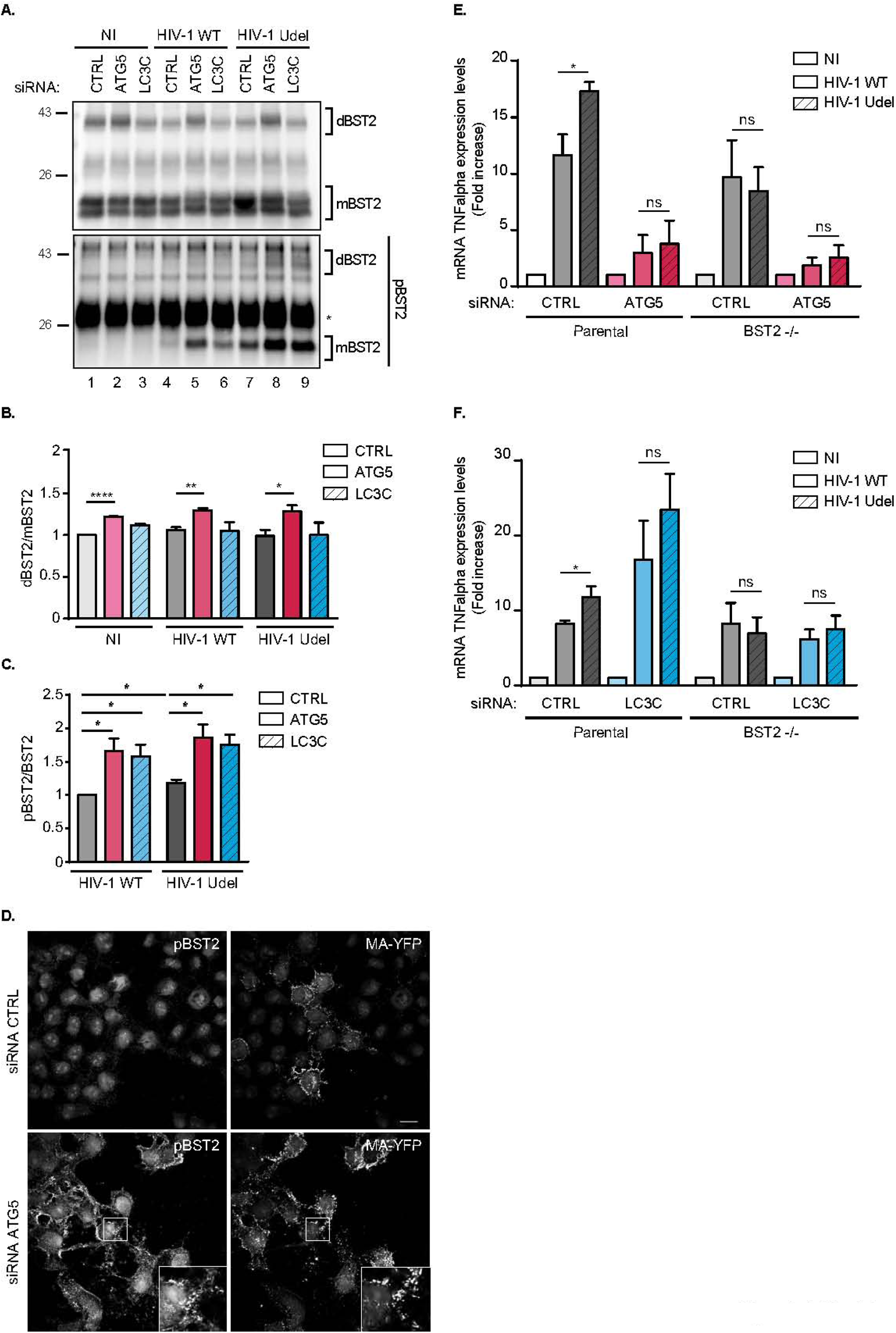
ATG5 selectively engages phosphorylated virus-tethered BST2 in an LC3 associated pathway controling Viral-induced BST2 signaling. **(A)** HeLa cells transfected with either control siRNA (CTRL) or siRNA targeting ATG5 or LC3C were infected with a VSV-G pseudotyped WT or Udel NL4.3 HIV-1 at a MOI of 0.5 for 24 hrs. Endogenous BST2 was immunoprecipitated with a mouse anti-BST2 antibody and deglycosylated precipitates were analyzed by western blot using rabbit anti-phosphorylated BST2 (pBST2) and rabbit anti-BST2 antibodies. **(B)** Quantification of the relative amount of the dimeric form of BST2 (dBST2) after normalization to the monomeric form (mBST2). Statistical analysis using two-tailed unpaired Student’s *t* test, mean ± SEM, n=3 experiments, * p ≤ 0.05, ** p ≤ 0.01, *** p ≤ 0.001. **(C)** Quantification of the relative amount of the phosphorylated monomeric form of BST2 (pBST2) after normalization to immunoprecipitated monomeric BST2. Statistical analysis using two-tailed unpaired Student’s *t* test, mean ± SEM, n=3 experiments, * p ≤ 0.05, *** p ≤ 0.001. dBST2: dimer of BST2; mBST2: monomer of BST2. **(D)** Confocal fluorescence microscopy of HeLa cells infected with VSV-G-pseudotyped WT HIV-1 MA-YFP for 24hrs following total staining with rabbit anti-pBST2 antibodies. Scale Bar: 20µM. All Western blots and immunofluorescence images presented are representative of at least three independent experiments. **(E)** HeLa BST2 WT and HeLa BST2 -/- cells transfected with either control siRNA (CTRL) or siRNA targeting ATG5 were transfected with the WT or Vpu-deleted (Udel) NL4.3HIV-1 provirus for 28 hrs. Total RNA was analyzed for TNFalpha mRNA levels relative to KDSR by qRT-PCR. Statistical analysis using two-tailed unpaired Student’s *t* test, mean ± SEM, n=3 experiments, * p ≤ 0.05. **(F)** HeLa BST2 WT and HeLa BST2 -/- cells transfected with either control siRNA (CTRL) or siRNA targeting LC3C were transfected with the provirus HIV-1 NL4.3 WT or Udel for 28 hrs. Total RNA was analyzed for TNFalpha mRNA levels relative to KDSR by qRT-PCR. Statistical analysis using two-tailed unpaired Student’s *t* test, mean ± SEM, n=3 experiments, * p ≤ 0.05.

Multiples studies correlate the multimerization state of BST2 upon viral tether and its phosphorylation on tyrosine residues present in its cytosolic domain (Y6 and Y8) (7, 21, 32). It has also been previously documented that the sensing function of BST2 is interrelated to its phosphorylation status and leads to the activation of the NFĸB pathway. We thus explored the phosphorylation status of the virus-tethered BST2 in cells depleted for ATG5. In accordance with previous studies (8), we confirmed an increase of the level of pBST2 upon infection, notably in cells infected with HIV-1 Udel (Figure 5A, lower panel, compare lanes 4-7 to lane 1, and figure 5C). Interestingly, depletion of ATG5 and LC3C strongly enriched the monomeric and dimeric forms of phosphorylated BST2 in cells infected with HIV-1 WT and Udel, compared to control cells (Figure 5A, lower panel, compare lanes 5-6 and 8-9 to 2-3, and figure 5C). Accordingly, confocal analysis of pBST2 staining revealed an accumulation of pBST2 in HIV-1 WT infected cells upon ATG5 depletion compared to control cells (Figure 5D). Furthermore, these patches of pBST2 were distributed in the vicinity of HIV-1 Gag product, MA/YFP at or near the plasma membrane (Figure 5D, lower panel, inset).

Altogether, these data support the view that ATG5 as well as LC3C might promote the degradation of phosphorylated forms of BST2 induced by virions retention through an LC3C-associated pathway

### LC3C-associated process regulates BST2-mediated signaling induced by virions retention

Phosphorylated BST2 tethering viruses have been regarded as signaling complexes that induced NFkB-dependent pro-inflammatory responses. We thus speculated that ATG5 as well as LC3C, via an LC3C-associated pathway, exert an effect on BST2-mediated signaling. To examine the involvement of these two ATG proteins, parental and BST2 knockout HeLa cells (BST2-/-) (34) were co-transfected with control (CTRL) siRNA or siRNA targeting ATG5 or LC3C and the WT (HIV-1 WT) or Vpu-defective (HIV-1 Udel) HIV-1 provirus. The outcome of these extinctions on the NFkB-dependent target gene expression TNFalpha was evaluated. To solely analyze the contribution of ATG5 and LC3C in the control of BST2-induced pro-inflammatory responses by Vpu, we normalized values obtained to the non-infected (NI) condition of each cell line. As previously reported (21), the expression of HIV-1 Udel provirus in parental control cells leads to a higher level of mRNA for the proinflammatory cytokine TNFalpha than those observed upon expression of HIV-1 WT provirus (Figures 5E and 5F). This increase was lost in BST2-/- cells, underlying the BST2 dependency of the enhanced proinflammatory response observed in these cell lines upon HIV-1 Udel expression (Figures 5E and 5F and Supplemental figure 5A). Surprisingly, the depletion of ATG5 decreased TNFalpha mRNA induction observed upon HIV-1 Udel provirus expression at a level similar to the one obtained with HIV-1 WT provirus expression (Figure 5E and Supplemental figure 5A), suggesting that the enhanced proinflammatory response observed upon HIV-1 Udel expression is dependent on both ATG5 and BST2 expression. Interestingly, knockdown of LC3C led to an increased level of TNFalpha mRNA upon expression of HIV-1 WT and Udel provirus in parental cells (Figure 5F and Supplemental figure 5B), which was completely lost in BST2-/- cells. This suggests that LC3C is implicated in the attenuation of BST2-induced proinflammatory gene expression mediated by Vpu upon HIV-1 infection.

Altogether, these results indicate that ATG5 is critical to initiate BST2-mediated signaling, while LC3C and Vpu attenuate the inflammatory response induced by BST2.

## Discussion

BST2 is one of the main restriction factors for mammalian enveloped viruses release. It considerably reduces viral dissemination by physically trapping *de novo* formed viral particles at the surface of infected cells. Studies demonstrate that BST2 forms clusters at the HIV-1 budding sites (35, 36) and that virus-tethered BST2 can be endocytosed and targeted for lysosomal degradation (4). Moreover, coupled with its viral tethering function, BST2 acts as a sensor of HIV-1 budding. By tethering viruses, BST2 activates the NF-kB signaling pathway leading to the expression of pro-inflammatory genes, hence establishing an anti-viral state of infected cells (21). To counteract this restriction, HIV-1, notably through the viral protein Vpu, engages multiples mechanisms (3, 16, 37). Among them, an LC3C-associated pathway is usurped by Vpu to favor the removal of virus-tethered BST2 from the budding site and their targeting into single membrane compartments to promote their degradation (27). Indeed, the attachment of LC3 proteins to the single membrane vesicles results in an increased acidification and degradation of its content (22).

LC3-associated pathways are initiated by the recognition of a surface receptor by immune complexes, exogenous particles, or pathogens (22, 25, 29). The binding of these ligands to their receptors triggers the invagination of the plasma membrane in a single membrane structure decorated by LC3. Several receptors have been reported to initiate this process, such as TLR1/2, TLR2/6, TLR4, TIM4 and FcR, resulting in recruitment of some, but not all, members of the autophagic machinery to a single membrane compartment (phagosome or endosome). These pathways are orchestrated by the concerted action of specific ATG proteins (e.g., LC3, ATG5 and Beclin-1), independently of the autophagy preinitiation complex (38). The work presented here reveals that ATG5 is a key determinant in (i) the recognition of plasma membrane virus-tethered BST2 and (ii) the initiation of the LC3C-associated process subverted by Vpu. Indeed, i) ATG5 and BST2 assemble as a complex, independently of Vpu and LC3C (Figure 1), ii) ATG5 takes in charge BST2 molecules tethering viral particles at the cell surface (Figure 2) and iii) binding of ATG5 to BST2 is strengthened by HIV-1 production (Figure 3A). Moreover, BST2 preferentially binds to the unconjugated form of ATG5 (figures 3A) and HIV-1 expression reduces the formation of ATG5-ATG12 conjugate, making free ATG5 available (Figure 3B). ATG5 preferentially interacts with dimerized BST2 and ATG5-ATG12 conjugation is dispensable for this interaction (Figure 3C). These results support a contribution of ATG5 in the selection at the cell surface and the engagement of virus-tethered BST2 molecules in an LC3C-associated pathway that does not require its conjugation to ATG12.

LC3-associated pathway has been initially described in phagocytic cells as LC3B-associated phagocytosis, leading to engulfment and degradation of exogenous cargos (pathogens, cells…)(22, 29). However, recent studies have revealed other pathways linked to LC3. An LC3-associated pathway has been connected to endocytic pathway with LC3B-associated endocytosis (LANDO) to favor the degradation of b-amyloid protein and the recycling of its receptor to the cell surface (23). The LC3C-endocytic-associated-pathway (LEAP), is another LC3-associated pathway that has been recently described (39). This pathway involves the targeting of cell surface receptors or plasma membrane derived proteins *via* the endocytic machinery to ATG9-containing vesicles and ultimately autophagosome. In contrast to the LAP or LANDO, involving the lipidation of LC3B, the LEAP pathway does not rely on direct LC3C lipidation to vesicular membranes, but instead on LC3C localization to endocytic compartment. While the stimuli, represented by the engagement of a plasma membrane receptor and the fate implying the recycling or the degradation of these engaged receptor are known, intracellular adaptors that activate these pathways is not yet known. In our study, we reveal that *via* its interaction with viral-tethered BST2, ATG5 engages an alternative route for the internalization of cell surface HIV-1 clusters, an LC3C-associated process, targeting them to degradation. Multiples studies show that ATG5 in its conjugated and unconjugated forms can modulate cellular signaling pathways in addition to its traditional role in autophagy (40). This unconventional role for ATG5 at the plasma membrane in the LC3C-associated process induced by HIV-1 infection has never been described and could open new avenue in the discovery of autophagy-independent functions of this protein, as an adaptor for receptor at the plasma membrane initiating an LC3-associated process.

We thus propose a model in which (1) the recognition of virus-tethered BST2 by ATG5 initiates the LC3C-associated pathway. (2) This engagement induces their endocytosis in a single membrane vesicle. (3) Then, through its interaction with the transmembrane domain of BST2, Vpu recruits selectively LC3C onto this vesicle membrane (27). (4) The presence of ATG5 at the level of this vesicle, *via* its interaction with BST2, will favor, after its conjugation with ATG12, the lipidation of LC3C and thus its anchoring to the vesicular membrane. Next, (5) the LC3C recruitment will accelerate the docking of this compartment to the lysosomes leading to the degradation of virus-tethered BST2 (Figure 6, left panel). Conversely, in absence of ATG5, the virus-tethered BST2 remain docked at the plasma membrane and virions accumulate in large clusters that are not endocytosed (Figure 6, right panel).

**Figure 6.**
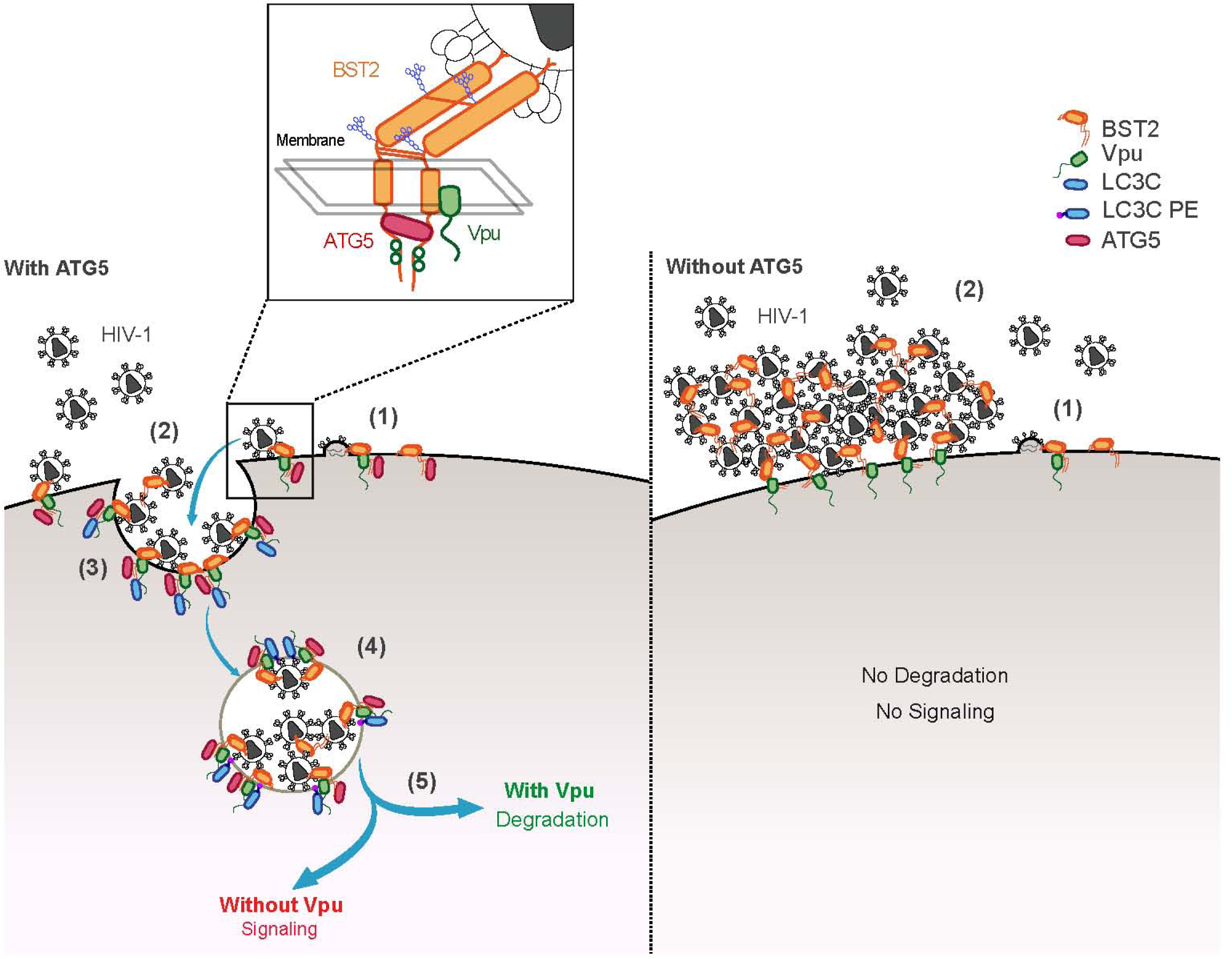
Model for the action of ATG5 in the selection and engagement of virus-tethered BST2 in Vpu-subverted LC3-associated pathway. **(Left panel)** (1) ATG5 initiated the LC3-associated pathway by recognizing and targeting phosphorylated virus-tethered BST2. (2) The interaction induces the endocytosis via regulators of clathrin-mediated vesicular trafficking of viral-tethered pBST2 in a single membrane compartment. (3) LC3C will be recruited to the BST2-ATG5 complex through its interaction with Vpu. (4) The presence of ATG5 at the level of the compartment will favor the lipidation of LC3C and thus its anchoring to this compartment sequestering virus-tethered pBST2. (5) The LC3C involvement will accelerate the docking of this compartment to the lysosomes leading to the degradation of virus-tethered pBST2. Upon Vpu expression, the degradation of virus-tethered pBST2 will attenuate the signaling mediated by BST2. Without Vpu expression, viral-tethered pBST2 will still be endocytosed. However, in absence of LC3C recruitment onto this compartment, the fusion with lysosome will be delayed and BST2-mediated signaling will persist. **(Right panel)** Without ATG5, virus-tethered BST2 will not be endocytosed and degraded. The virion retention mediated by BST2 will not induce the activation of NF-ĸB signaling.

Our model hypothesizes that ATG5 engages specific forms of BST2 tethered to virions to an LC3C-associated pathway, in which endosomal structures are customized by some ATG proteins. Interestingly, recent proteomic approaches described strong link between ATG5 and clathrin adaptor protein complexes (AP)-dependent endocytic pathway (41). Moreover, several studies reported that Vpu/BST2 complex can interact with the AP complexes and that these interactions are important for BST2 antagonism by Vpu (11). It is thus tempting to speculate that ATG5 and regulators of clathrin-mediated vesicular trafficking acts in concert to target phosphorylated and dimerized forms of BST2 tethering virions for the degradation in an LC3C-associated pathway.

Indeed, analysis of ATG5-BST2 interaction showed that ATG5 selectively engages phosphorylated BST2 (pBST2) in the LC3C-associated pathway induced by HIV-1 infection. ATG5 depletion leads to a marked accumulation of pBST2 (Figures 5A and 5C), implying that BST2 acts like a sensor for the LC3C-associated pathway. Multiples studies showed that the viral sensing function of BST2, after phosphorylation of its tyrosine residues Y_6_Y_8_, induces the activation of NFĸB signaling (7, 8, 21, 32). This induction of the NFkB signaling pathway requires the internalization of BST2 (7) and Vpu is able to attenuate BST2-induced pro-inflammatory responses (21). LC3-associated pathways as well as the membrane trafficking pathway, in regulating transport of internalized receptors, are known to be an important signaling regulatory network. Here, we found that BST2-mediated signaling induced by HIV-1 infection is impaired in ATG5-depleted cells in HeLa, whereas this signaling is increased in LC3C-knockdown cells (Figures 5E and F). These data support the view that the LC3C-associated pathway subverted by Vpu is implicated in the control of the inflammatory response mediated by BST2. Therefore, to complement our model, we propose that the recruitment by ATG5 of the phosphorylated virus-tethered BST2 at the plasma membrane and the engagement of these complexes onto an LC3C-associated pathway, can additionally lead to the activation of NF-ĸB signaling (Figure 6). Next, Vpu, by recruiting LC3C on the vesicles that sequestered virus-tethered pBST2, enhances the degradation of these complexes and thus attenuates the proinflammatory response mediated by viral tethering (Figure 6, left panel, compare the model with and without Vpu). Without ATG5, virion retention mediated by BST2 does not trigger the activation of NF-ĸB signaling (Figure 6, right panel).

Previous report showed that Vpu is able to inhibit NFkB activation *via* the sequestration of b-TrCP, an E3 ubiquitin ligase responsible for the ubiquitination and degradation of IkB (7, 42). Interestingly, Sauter et al. reported that Vpu’s inhibitory effect on BST2-mediated NFkB signaling could not be solely account for its interaction with b-TrCP (43). We can thus postulate that the subversion of the LC3C-associated pathway by Vpu is the part of the mechanism by which Vpu attenuates BST2-mediated NFKB activation. The Vpu-mediated antagonism of BST2 restriction is a multifaceted process involving so far, the AP1/2-mediated mistrafficking, the b-TRCP-mediated ubiquitination and the ESCRT machinery. The discovery of an LC3-associated process implicated in the targeting of virus-tethered BST2 from the viral budding site to the degradation adds another level of complexity and thus how these pathways might work together need to be elucidated.

BST2 has a broad activity against diverse families of enveloped viruses (44). Many lipid-enveloped viruses could be restricted by BST2 (45). Therefore, further studies will required to elucidate whether the LC3-associated pathway could be subverted by other viruses to counteract BST2 restriction and/or attenuate the cellular anti-viral state established by BST2.

## Materials and Methods

### Cell lines

HeLa and HEK293T cells were grown in DMEM (Dulbecco’s modified Eagle’s medium) with GlutaMAX and 10% decomplemented-FCS (fetal calf serum) (Gibco, Life Technologies). HeLa WT and HeLa BST2 KO cells were a kind gift of Stéphane Fremont (Institut Pasteur, France).

### CRISPR-Cas9 knockout in HeLa cells

Expression plasmids for single guide RNA (sgRNA) (pMLM3636 vector, a gift from Keith Joung; Addgene plasmid #43860) targeting an exon within ATG5 genes were transiently transfected into HeLa cells using Lipofectamine LTX with PLUS Reagent (Life technologies), along with a plasmid expressing Cas9 fused with GFP (pCas9 GFP, gift from Kiran Musunuru: Addgene plasmid #44719). The following target sequences were used: 5’-AACTTGTTTCACGCTATATC-3’ (ATG5, exon 1; guide 129) and 5’- AAGATGTGCTTCGAGATGTG-3’ (ATG5, exon 1; guide 92). 48 hours after transfection, GFP expressing HeLa cells were sorted with BD FACS Aria III Cell Sorter, cultivated for 7 days in complete medium, cloned by limiting dilution in 96-well flat-bottomed culture plates and expanded for 15 days. The clones were then screened by PCR amplification of the targeted region of the genome. The PCR products were cloned into pCR-Blunt II TOPO (Life technologies) and up to 20 independent clones were sequences in each case. The primer pair used for PCR amplification of the sgRNA target sites was: ATG5fw 5’-TCCAAAATAAGCATGAATTAGCTGT-3’ and ATG5rev 5’- TGGGCTTGAAAGACTGATGCA-3’ (ATG5 target site). ATG5 knockout was checked by western blot. CRISPR LC3C cell lines were previously described (27).

### Small interfering RNA and transfection

Cells were transfected with relevant small interfering RNA (siRNA) oligonucleotide using Lipofectamine RNAiMAX (Life Technologies), according to the reverse transfection procedure described in the manufacturer’s recommendations. The final concentration of siRNA oligonucleotides was 7.5 to 30 nM. 21-nucleotide RNA duplexes with 2-nucleotide 3-(2-deoxy) thymidine overhangs were used. Specific siRNA sequence targeting ATG5 was used: ATG5 (5’-GGATGCAATTGAAGCTCATdTdT-3’, positions 647-665 for variant 1, 574-592 for variant 2, 519-537 for variant 3, 647-665 for variant 4, 412-430 for variant 6 and 284-302 for variant 7). The siRNA targeting LC3C (5’- GCTTGGCAATCAGACAAGAGGAAGT-3’, position 143-167) was previously described (27, 46). The On-Target plus SMART pool siRNA targeting BST2 (L-011817-00) was purchased from Dharmacon. The siRNA (D-001810-01 from Dharmacon) was used as negative control (referred as siRNA Control).

### Mammalian expression vectors and transfection

NL4-3 HIV-1 proviral DNA were obtained, respectively from NIH AIDS Research and Reference Reagent Program (Division of AIDS, NIAID). NL4-3 Udel HIV-1 proviral DNA and wild type NL4-3 (MA/YFP) HIV-1 was, respectively, a gift from Dr. K. Strebel (47) and Dr. P. Bienasz (48). HIV-1 NL4-3 Vpu ORF was cloned into pEGFP-N1 (Clontech, France). The ORF of human ATG5 was cloned in frame with HA tag into pAS1B vector (pAS1B-HA-ATG5). The ORFs of human LC3C, human ATG5 and human Beclin 1 were cloned in frame with FLAG or HA affinity tag into the p3XFlag vector (p3XFlag-LC3C, p3XFlag-ATG5 and p3XFlag-Beclin 1) or pCMV-HA (pCMV-HA-LC3C). The ORFs of human ATG5 was cloned in frame with GFP tag into the pEGFP-C1 vector. The cDNA encoded the WT and short isoforms of human BST2 was cloned in pcDNA3.1 vector (pcDNA-BST2 WT; pcDNA-BST2 M1A) and in frame with FLAG tag in p3XFlag vector (p3XFlag-BST2 WT). The mutated forms of BST2 (BST2 2-4-AAA, BST2 3-5-AAA, BST2 4-6-AAA, BST2 5-7-AAA, BST2 6-8-AAA, BST2 7-9-AAA, BST2 8-10-AAA, BST2 9-11-AAA, BST2 10-12-AAA, BST2 11-13-AAA, BST2 12-14-AAA, BST2 13-15-AAA, BST2 14-16-AAA, BST2 15-17-AAA, BST2 16-18-AAA, BST2 17-19-AAA, BST2 18-20-AAA, BST2 19-21-AAA, BST2 Y_6_Y_8_-AA, BST2 C3A, BST2 M1A) and ATG5 (ATG5-K130R) were made by PCR mutagenesis using the QuikChange II XL site directed mutagenesis kit (Stratagene, France). The cDNA encoded ATG5-K130R was cloned in frame with GFP tag into the pEGFP-C1 vector. Similarly, the deleted form of BST2 (BST2 Delta CT) was generated by site directed mutagenesis by introducing a stop codon in position 21 of BST2 in p3XFlag-BST2 WT. Mutagenesis and subclonings were verified by DNA sequencing.

Transfections of HeLa cells or HEK293T cells with mammalian expression vectors were performed using Lipofectamine LTX with PLUS Reagent (Life technologies), following the manufacturer’s instructions.

### Viral stocks

Stocks of VSV-G pseudotyped wild type (WT) NL4-3 HIV-1, NL4-3 Udel HIV-1, WT NL4-3 (MA/YFP) HIV-1 and Udel NL4-3 (MA/YFP) HIV-1 were obtained by transfection of HEK293T cells with HIV-1 proviral DNA along with a VSV-G expression vector (pMD.G) and polyethylenimine (PEI) (Polysciences). Twenty-four hours after transfection, cells media were removed and cells were cultured for additional 24hrs in fresh media. Supernatants were then collected and filtered (0.45 µm). Viral titers were determined by infection of HeLa cells with serial dilutions of the viral stocks for 24hrs, followed by flow cytometry analysis of CAp24 antigen expression on fixed and permeabilized cells labelled with KC57-fluorescein isothiocyanate (FITC) (Beckman Coulter).

### Antibodies

The following antibodies were used for immunoblotting and/or immunofluorescence: Mouse monoclonal anti-CAp24 HIV-1 (National Institute for Biological Standards and Control Centralized Facility for AIDS Reagents (NIBCS); ARP366), human monoclonal anti-SUgp120 (National Institute for Biological Standards and Control Centralized Facility for AIDS Reagents (NIBCS); 2G12), mouse monoclonal anti-BST2 (Abnova; H00000684-M15), mouse monoclonal anti-GAPDH (Santa Cruz; sc-47724), mouse monoclonal anti-α-Tubulin (Sigma-Aldrich; T9026), mouse monoclonal anti-β-Actin (Sigma-Aldrich; A2228), mouse monoclonal anti-Flag (M2)-HRP (Sigma-Aldrich; A8592), rabbit polyclonal anti-ATG5 (Cell Signaling Technology; 12994S), rabbit polyclonal anti-LC3B (Novus Biologicals; NB600-1384), rabbit polyclonal anti-BST2 (NIH AIDS Reagent Program, Division of AIDS, NIAID, NIH; 11721) and rabbit polyclonal anti-Vpu (NIH AIDS Reagent Program, Division of AIDS, NIAID, NIH; 969). rat monoclonal anti-HA (3F10)-HRP (Roche; 12013819001), goat polyclonal anti-GFP-HRP (GeneTex; GTX26663).

Anti-phosphoBST2 antiserum was elicited in rabbits by using a modified peptide of BST2 (amino acids 1 to 21) phosphorylated on tyrosines in position 6 and 8, generating a polyclonal antibody against the tyrosine-phosphorylated intracellular portion of BST2 (Proteogenix).

Secondary antibodies against the mouse, rabbit, goat or human immunoglobulin G coupled to Alexafluor-594, Alexafluor-488 or Alexafluor-647 (purchased from Invitrogen) were used for immunofluorescence. Secondary antibodies against the mouse and the rabbit immunoglobulin G coupled to HRP (Dako) were used for immunoblotting experiments.

The antibodies used for flow cytometry are Alexa Fluor 647 mouse anti-human BST2 (Biolegend; 348404) and HIV-1 core antigen-FITC (KC57, Beckman Coulter; 6604665).

### Immunoprecipitation assay

HEK293T or HeLa cells were co-transfected with mammalian expression vectors and HIV-1 proviral DNA NL4-3 (WT or Udel) as described above. Twenty-four hours after transfection, immunoprecipitation of the relevant protein was performed.

HeLa cells were treated with siRNA (7.5-30 nM) as described above. Forty-eight hours after transfection, siRNA-treated cells were infected with VSV-G pseudotyped NL4-3 (WT or Udel) HIV-1 for 2h30 at a multiplicity of infection (M.O.I.) of 0.5. Twenty-four hours after transfection, immunoprecipitation of the relevant protein was performed. To observed pBST2 after immunoprecipitation, cells were incubated for 5 min in DMEM containing 100uM of pervanadate before lysis.

Cells were lysed in ice-cold lysis buffer (50 mM Tris HCl pH 7.4, 150 mM NaCl, 0.1% SDS, 0.5% sodium deoxycholate, 1% NP-40, 200 µM sodium orthovanadate) with complete protease inhibitor cocktail (Roche). The protein concentrations were determined using a Bradford protein assay (Bio-Rad), and equal amounts of protein for each sample were used for the following steps. Immunoprecipitations were performed by incubating indicated whole cell extracts overnight at 4°C with monoclonal mouse anti-Flag or monoclonal mouse anti-BST2 antibody, or mouse IgG CTRL coupled to Dynabeads protein G (Life Technologies). The beads were washed 4 times with lysis buffer, and proteins were eluted in 2X Laemmli (Sigma-Aldrich) or deglycosylated by an 1h treatment with PNGase F (NEB) then eluted in 2X Laemmli as previously described.

GFP-tagged proteins were immunoprecipitated using GFP-TRAP beads (Chromotek) using TNTE buffer (20mM Tris-HCL pH 7.4, 150mM NaCl, 5mM EDTA, 0.5% Triton X-100) with complete protease inhibitor cocktail. Immunoprecipitations were performed by incubating indicated whole cell extracts for 1h30 at 4°C with GFP-TRAP beads. The beads were washed and proteins were eluted as previously described.

### Western Blotting

Cell lysates were lysed in ice-cold DOC buffer (10mM Tris, pH 8, 150mM NaCl, 1mM EDTA, 1% Triton X-100 and 0.1% DOC 10%) containing complete protease inhibitor cocktail (Roche). Lysates were cleared by centrifugation for 15 min at 13,000 rpm. The protein concentrations were determined using a Bradford protein assay (Bio-Rad), and equal amounts of protein for each sample were used for the following steps Cell lysates and immunoprecipitated proteins were subjected to SDS-PAGE gels. Laemmli 2x concentrate (Sigma) has been used as a sample buffer for reducing and loading protein samples in SDS-PAGE, whereas NuPAGE LDS sample buffer 4x concentrate (ThermoFisher Scientific) has been used for non-reducing conditions. Proteins were then transferred onto hydrophobic polyvinylidene difluoride membranes (PVDF, 0.45 µm, Millipore), followed by blocking in milk buffer (Tris-buffered saline [TBS] [0.5 M Tris pH 8.4, 9% {wt/vol} NaCl], 5% [wt/vol] nonfat dry milk, 0.05% [vol/vol] Tween 20) for 1h at room temperature (RT). Membranes were incubated overnight at 4°C with the appropriated primary antibodies in milk buffer or BSA buffer (Tris-buffered saline [TBS] [0.5 M Tris pH 8.4, 9% {wt/vol} NaCl], 3% [wt/vol] BSA, 0.05% [vol/vol] Tween 20). Blots were washed with TBS containing 0.05% (vol/vol) Tween 20 and incubated with appropriate HRP-conjugated secondary antibodies in milk or BSA buffer for 1h at RT. After washing, protein bands were detected by using Amersham ECL Select Western blotting detection reagent (GE Healthcare).

### Formaldehyde crosslinking of BST2

To stabilize non-covalent oligomers of BST2 for SDS-PAGE analysis and immunoblotting, cells were harvested using PBS-EDTA 1mM, pelleted, washed once with PBS then resuspended in 1ml of serum-free DMEM containing 1% formaldehyde. Cells were incubated at 37°C for 20min under rotation. The reaction was stopped by adding 100µl of 1.25M glycine in PBS and cells were incubated 5 min at RT under rotation. Cells were then pelleted and lysed in DOC buffer. Lines were drawn on each BST2 signal and resulting line plots of the chemiluminescence intensity were used to confirm the enrichment of N-linked glycosylated form of BST2.

### Quantitative RT-PCR

Total cellular RNA was extracted using the Reliaprep RNA Cell Miniprep System kit (Promega; Z6012) following the manufacturer’s instructions. For each sample, 500ng to 2µg of total RNA were subjected to DNase I treatment (TURBO DNase; ThermoFisher; cat AM2239) and cDNA synthesis was performed using the High-capacity cDNA Reverse Transcription Kit (Applied Biosystems; 4368814). The different mRNA levels were assayed using SYBR Green Supermix (Bio-Rad; 1725275) in a real-Time PCR detection system (LightCycler® 480). The PCR conditions and cycles were as follows: an initial DNA denaturation at 95°C for 5 min, followed by 45 cycles of amplification (denaturation: 95°C for 10 sec, annealing: 63°C for 10 sec and extension :72°C for 10 sec), followed by a melting-curve analysis cycle. Each point was performed in technical duplicate. The relative abundance of TNFα mRNA (sense: 5’- TCCTTCAGACACCCTCAACC-3’ ; antisense: 5’-AGGCCCCAGTTTGAATTCTT-3’) was calculated by the comparative ΔΔCt method normalizing to the housekeeping product KDSR mRNA (sense : 5’- AGATGAGTTGGACCCATTGC -3’ ; antisense : 5’-AAGCCATGAGTTTCCACCAG -3’) in HeLa cells.

### Fluorescence microscopy

Cells were grown on coverslips, transfected with siRNA, infected, and then fixed with 4% paraformaldehyde in PBS for 20 min. For staining of pBST2, cells were incubated for 5 min in DMEM containing 100uM of pervanadate before fixation with 4% paraformaldehyde. Cells were then washed three times using PBS containing 50mM NH_4_Cl. Cells were permeabilized with 0.2% Triton X-100 for 10 min. Coverslips were blocked with 5% BSA for 1h, incubated with primary antibody in 1% BSA for 1h, washed with PBS, and incubated with secondary antibody in 1% BSA for 30 min, before final washing with PBS and MilliQ water.

For extracellular staining of BST2, living cells were incubated for 1h at 4°C with rabbit polyclonal anti-BST2 (NIH) together with mouse anti-Env SUgp120 (110H, Hybridolab). Then, cells were fixed with 4% PFA in PBS for 20 min and labelled with appropriate fluorophore-conjugated secondary antibodies. Cells were mounted in DAPI Fluoromount-G (SouthernBiotech). Microscope IXplore spinning disk Olympus was used for confocal analysis. We used the 60X plan-apochromat objective, with a numerical aperture 1.42. Images were processed using ImageJ software. The experiments were repeated as indicated in the figure legends, and representative images are shown. In all experiments, images shown in individual panels were acquired using identical exposure times or scan settings and adjusted identically for brightness and contrast using Photoshop CS5 (Adobe).

### Electron Microscopy

The method used is essentially those described previously (16, 49). Briefly, ultrathin cryosections (50 nm) were stained with mouse antibodies against HIV-1 p24/p55 (EVA365 and EVA366, NIBSC), rabbit anti-mouse bridging antibody (Rockland Immunochemicals Inc. Limerick, PA), and 10 nm PAG (Protein A gold reagents were obtained from the EM Lab, Utrecht University, Utrecht, The Netherlands). Sections were fixed in 1% (v/v) glutaraldehyde for 10 min, embedded in uranyl acetate in methylcellulose, as described previously, and examined with a Technai G2 Spirit transmission electron microscope (FEI Company UK. Ltd., Cambridge, UK).

### Flow cytometry

Twenty-four hours post-infection, siRNA transfected HeLa cells were washed twice in PBS and collected using PBS EDTA 1mM. Cells were first stained with LIVE/DEAD™ Fixable Violet Dead Cell Stain Kit (Thermofisher) according to the manufacturer’s recommendations. Then, cells were washed in cold PBS/1% (w/v) BSA and stained for 1 hour at 4°C with AlexaFluor647-conjugated anti-BST2 (Biolegend) or control isotype (Biolegend). The cells were washed three times in cold PBS/1% (w/v) BSA, then fixed in 4% paraformaldehyde (PFA) and permeabilized in PBS/1% BSA/0.1% saponin before staining with a FITC-conjugated anti-Cap24 (KC57-FITC, Beckman Coulter, France) for 1h at room temperature. Cells were washed and analysed using the BD LSRFortessa™ cell analyzer.

### HIV-1 production assay

For a HIV-1 production assay, HeLa BST2 WT and HeLa BST2 KO cells were treated with siRNA (7.5-30 nM) as described above and transfected with HIV-1 proviral DNA NL4-3 (WT or Udel). Twenty-eight hours after transfection, cell lysates were analyzed by qRT-PCR and western blotting.

For CD4+ T cells infection, CRISPR-Cas9 knockout cells were activated for three days as described previously, then infected by spinoculation with VSV-G pseudotyped NL4-3 (WT or Udel) HIV-1 for 2h at a multiplicity of infection (M.O.I.) of 0.5. Forty-eight hours after infection, cell lysates were analyzed by qRT-PCR and western blotting.

In a single round of infection, HeLa cells were treated with siRNA (7.5-30 nM) as described above. Forty-eight hours after, siRNA-treated Hela cells were infected with VSV-G pseudotyped NL4-3 (WT or Udel) HIV-1 for 2h30 at a multiplicity of infection (M.O.I.) of 0.5. Thirty-two hours after infection, media was removed and replaced with fresh media for additional 16hrs. Supernatants were then collected, 0.45 µm-filtered and used for HIV-1 CAp24 quantification by ELISA (released CAp24) (Perkin Elmer). Cell lysates were analyzed by western blotting.

### Statistical analysis

The statistical details of all experiments are reported in the figure including statistical analysis performed, error bars, statistical significance, and exact n numbers. Statistics were performed using GraphPad Prism 6 software, as detailed in the figure legends.

## Acknowledgements

We thank Stéphane Emiliani, Florence Margottin-Goguet and Mark Scott for helpful discussions and for critical reading of the manuscript, Annegret Pelchen-Mathews for IEM images. We thank all the members of the “Interactions hôte-virus” Laboratory for comments and helpful discussions. We thank the Imaging photonic Facility IMAG’IC and the Cytometry and Immunobiology Facility CYBIO of the Cochin Institute for technical assistance. The following reagents were obtained through the National Institute for Biological Standards and Control Centralised Facility for AIDS Reagents, which is supported by the EU Program EVA and the UK Medical Research Council: HIV-1 gp120 monoclonal antibody (2G12) from Dr H. Katinger and mouse antibodies against CAp24 (EVA365) from B. Wahren. The following reagents were obtained through the AIDS research and reference reagent program, Division of AIDS, NIAID, NIH: anti-human BST2 from Drs K. Strebel and A. Andrew, HIV-1 NL4-3 Vpu antiserum from Drs K. Strebel and F. Maldarelli and pNL4-3 from Dr. M. Martin. We thank Dr K. Strebel for the gift of pNL4-3/Udel proviral DNA and Dr. P. Bieniasz for the gift of pNL4-3(MA/YFP) proviral DNA. We thank Dr. S. Frémont for the gift of BST2 KO cell line. D.J. holds a fellowship from ANRS and then from SIDACTION, A.K., F.B. and P.V. from ANRS. M.P and L.L. hold a fellowship from the “Ministère Français de l’enseignement supérieur et de la Recherche”. This work is funded by ANRS and SIDACTION.

## Author contribution

DJ and CBT conceived the project; DJ, MV, FB, AK and CBT designed experiments; DJ, MV, FB, MP, PV, AK, LL and CBT performed the experiments; DJ, MV, FB, PV, AK and CBT analyzed and interpreted data; DJ and CBT wrote the manuscript with input from all listed authors; KJ, SGM, PV, DJ and CBT edited the manuscript.

## Supplemental data

**Supplemental figure 1.**
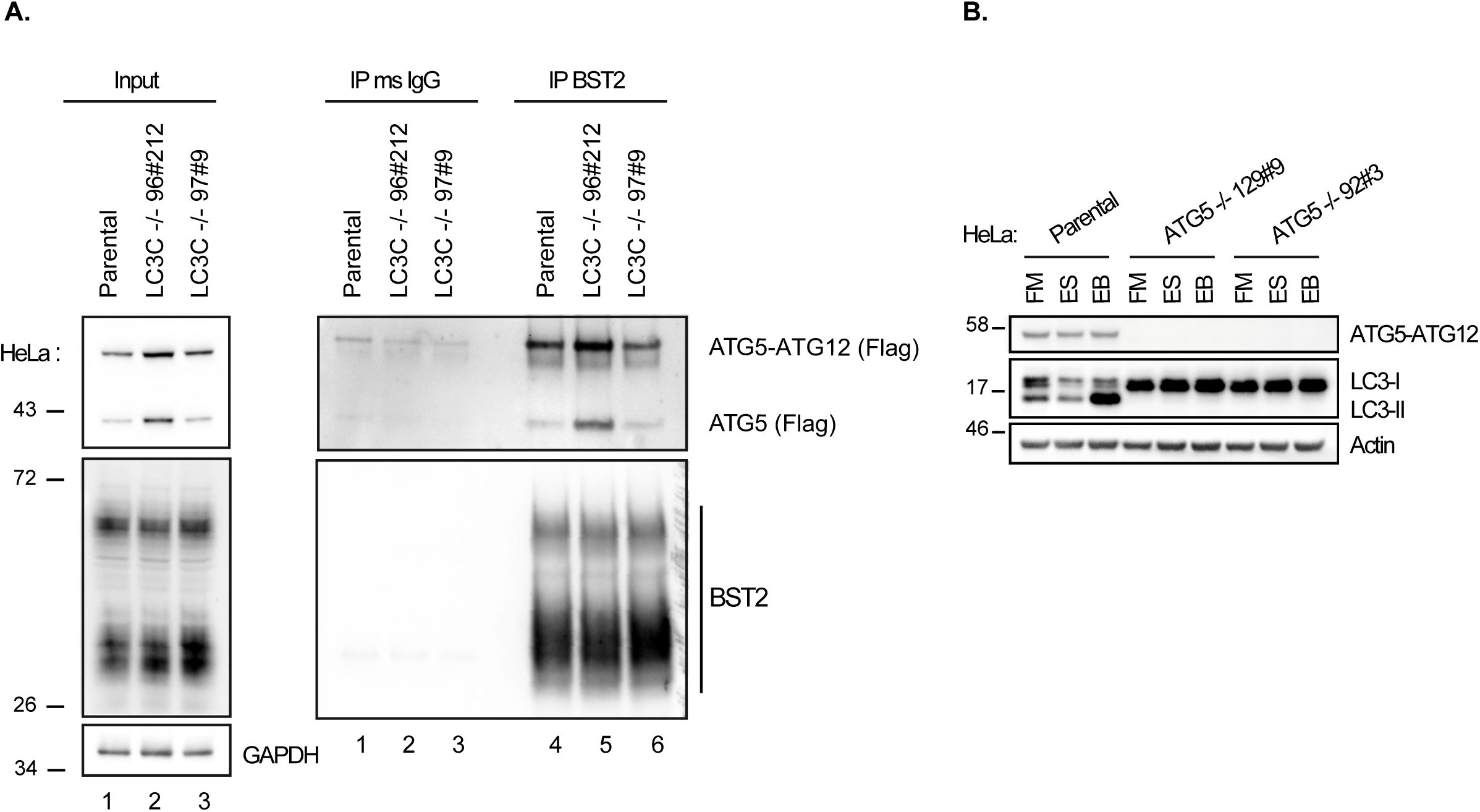
related to Figure 1. **(A)** Immunoprecipitation of endogenous BST2 in Parental (CTRL) or LC3C knockout (LC3C -/- 196#212 or 97#9) HeLa cells extracts co-transfected with p3XFlag-ATG5. Immunoprecipitated proteins were detected by western blotting using anti-BST2 and anti-Flag-HRP antibodies. **(B)** Parental (CTRL) or ATG5 knockout (ATG5 -/- 129#9 or 92#3) HeLa cells were incubated in full medium (FM) or EBSS for amino acid depletion (ES) without or with Bafilomycin A1 (EB) for 2 h before immunoblotting for ATG5, Actin, and LC3B

**Supplemental figure 2.**
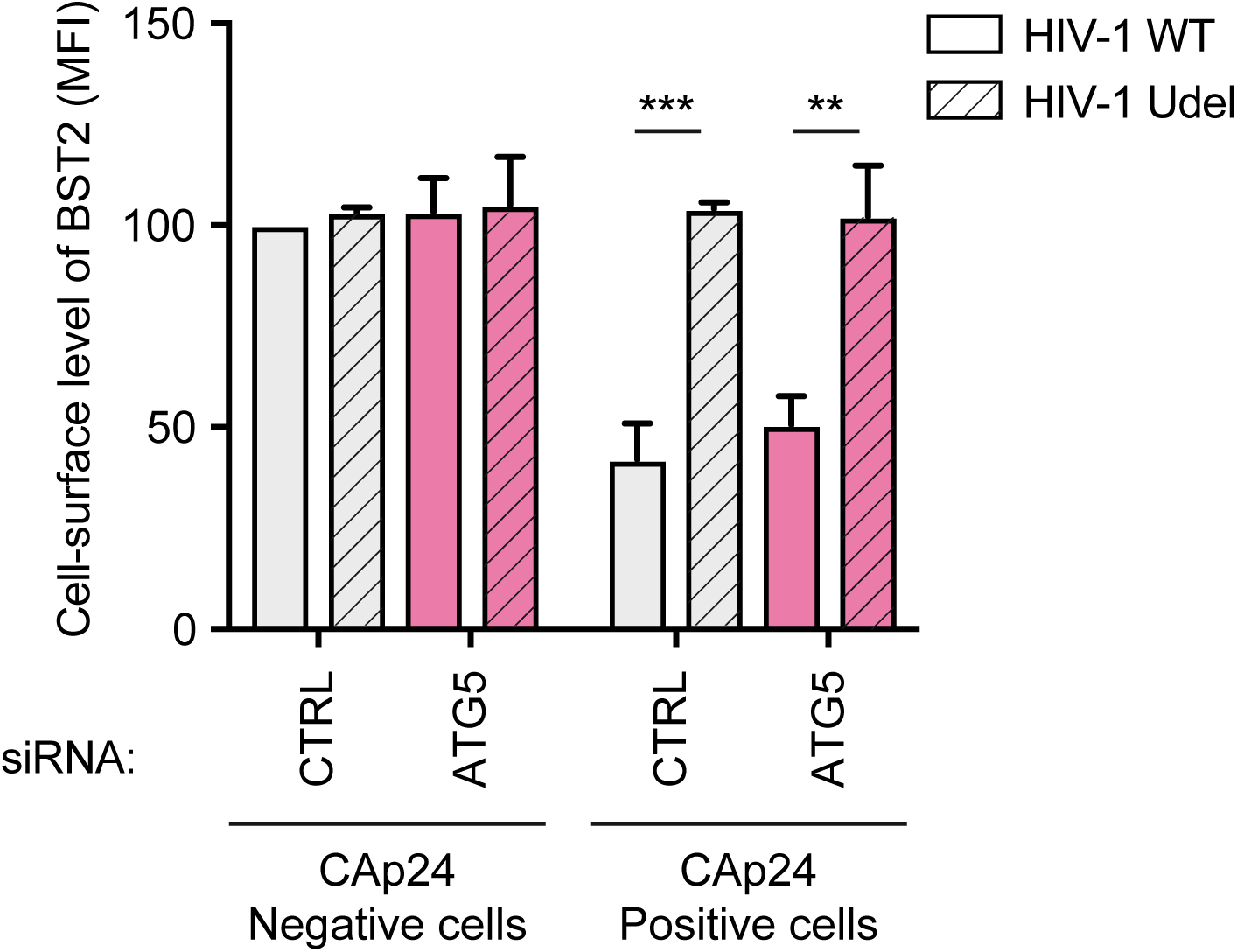
related to Figure 2. HeLa cells transfected with either control siRNA (CTRL) or siRNA targeting ATG5 were infected with a VSV-G pseudotyped WT or Udel NL4.3 HIV-1 at a MOI of 0.5. Twenty-four hours later, cells were stained at the cell surface with anti-BST2 antibody. Cells were then fixed, permeabilized and stained for Gag using anti-CAp24 antibody. Cells were then processed for flow cytometry analysis. Bar graphs represent cell surface level of BST2 in CAp24 negative and positive cells for each siRNA condition. Values are expressed as the Mean Fluorescence Intensity (MFI). Statistical analysis using two-way ANOVA with Tukey’s multiple comparison test, mean ± SEM, n=3 experiments; ** p ≤ 0.01, *** p ≤ 0.001.

**Supplemental figure 3.**
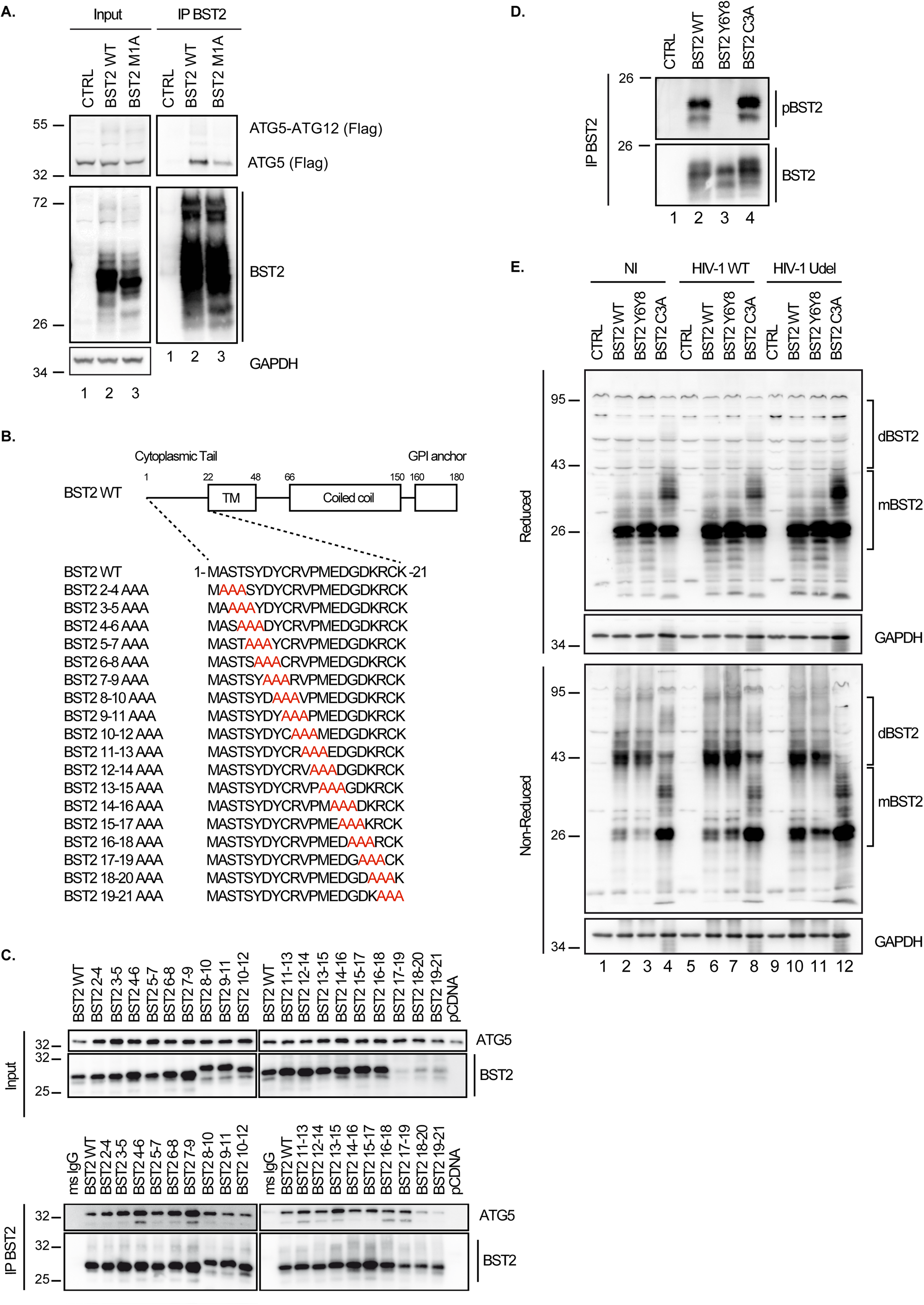
related to Figure 4. **(A)** HEK293T cells were transfected with p3XFlag-ATG5 and either pcDNA, pcDNA-BST2 WT, or pcDNA-BST2 M1A encoding for the short isoform of BST2. BST2 was immunoprecipitated with mouse anti-BST2 antibodies. Immunoprecipitated proteins were detected by western blotting using anti-flag-HRP and rabbit anti-BST2 antibodies. **(B)** Schematic representation of alanine mutagenesis in the cytoplasmic domain of BST2. **(C)** HEK293T cells were transfected with pCMV-HA-ATG5 and either pcDNA, pcDNA-BST2 WT, or plasmids encoding for mutated BST2 as described in (B). BST2 was immunoprecipitated with mouse anti-BST2 antibodies. Immunoprecipitated proteins were detected by western blotting using anti-HA-HRP and rabbit anti-BST2 antibodies. **(D-E)** HEK293T cells were co-transfected with either pcDNA-BST2 WT or BST2 Y_6_Y_8_ (double tyrosine phosphorylation defective mutant) or BST2 C3A (triple cysteine dimerization defective mutant) and the WT or Udel NL4.3 HIV-1 provirus. BST2 dimerization and phosphorylation were analyzed. For the detection of BST2 phosphorylation, BST2 was immunoprecipitated with a mouse anti-BST2 antibody and deglycosylated precipitates were analyzed by western blot using a rabbit antibody specific of phosphorylated tyrosine 6 and 8 of BST2 and a mouse anti-BST2 antibody (D). For BST2 dimerization, cell lysates were prepared under reducing or non-reducing conditions. dBST2: dimer of BST2; mBST2: monomer of BST2 (E). All Western blots presented are representative of at least three independent experiments.

**Supplemental figure 4.**
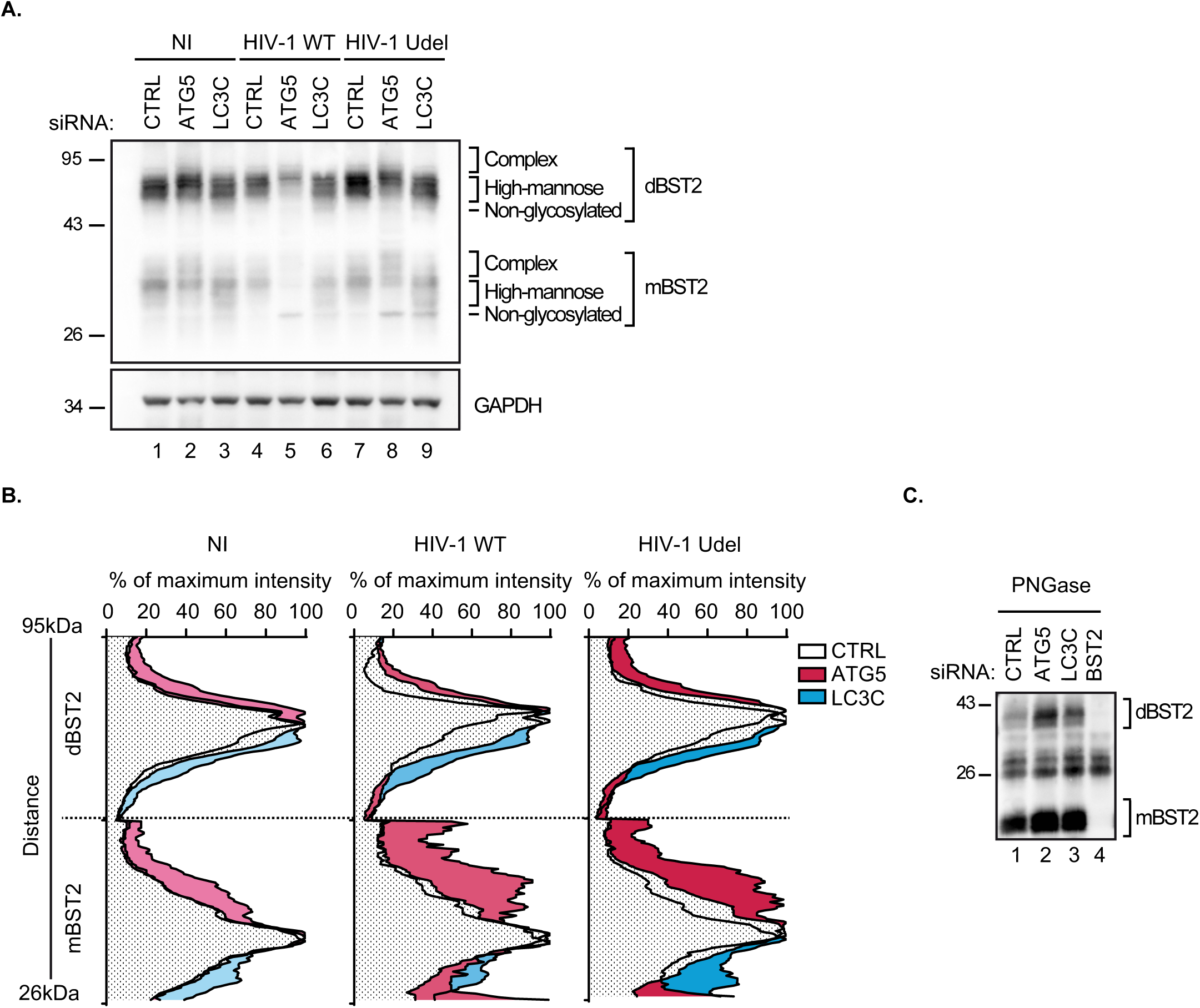
related to Figure 5. **(A)** HeLa cells transfected with either control siRNA (CTRL) or siRNA targeting ATG5 or LC3C were infected with a VSV-G pseudotyped WT or Udel NL4.3 HIV-1 at a MOI of 0.5 for 24 hrs. Cells were treated for 20min with formaldehyde 1% before preparing cell extracts under reducing conditions. Western blot analysis of BST2 and GAPDH in non-infected and infected siRNA-treated cells. **(B)** Line-scan profiles of BST2 intensity obtained on western blot of extracts of siRNA-treated HeLa cells infected with a VSV-G pseudotyped HIV-1 NL4.3 WT or Udel at a MOI of 0.5 for 24hrs and formaldehyde crosslinked. Analysis of BST2 patterns was determined by western blot and line-scan profiles of BST2 intensity were plotted across 95 to 26 kDa. The x-axis represents the % of maximum intensity for the monomeric and dimeric forms of BST2 and the y-axis, the distance between molecular weights. **(C)** HeLa cells transfected with either control siRNA (CTRL) or siRNA targeting ATG5, LC3C or BST2 were infected with a VSV-G pseudotyped WT or Udel NL4.3 HIV-1 at a MOI of 0.5 for 24hrs. Endogenous BST2 was immunoprecipitated with a mouse anti-BST2 antibody and deglycosylated precipitates were analyzed by western blot using a rabbit anti-BST2 antibody. dBST2: dimer of BST2; mBST2: monomer of BST2. All Western blots presented are representative of at least three independent experiments.

**Supplemental figure 5.**
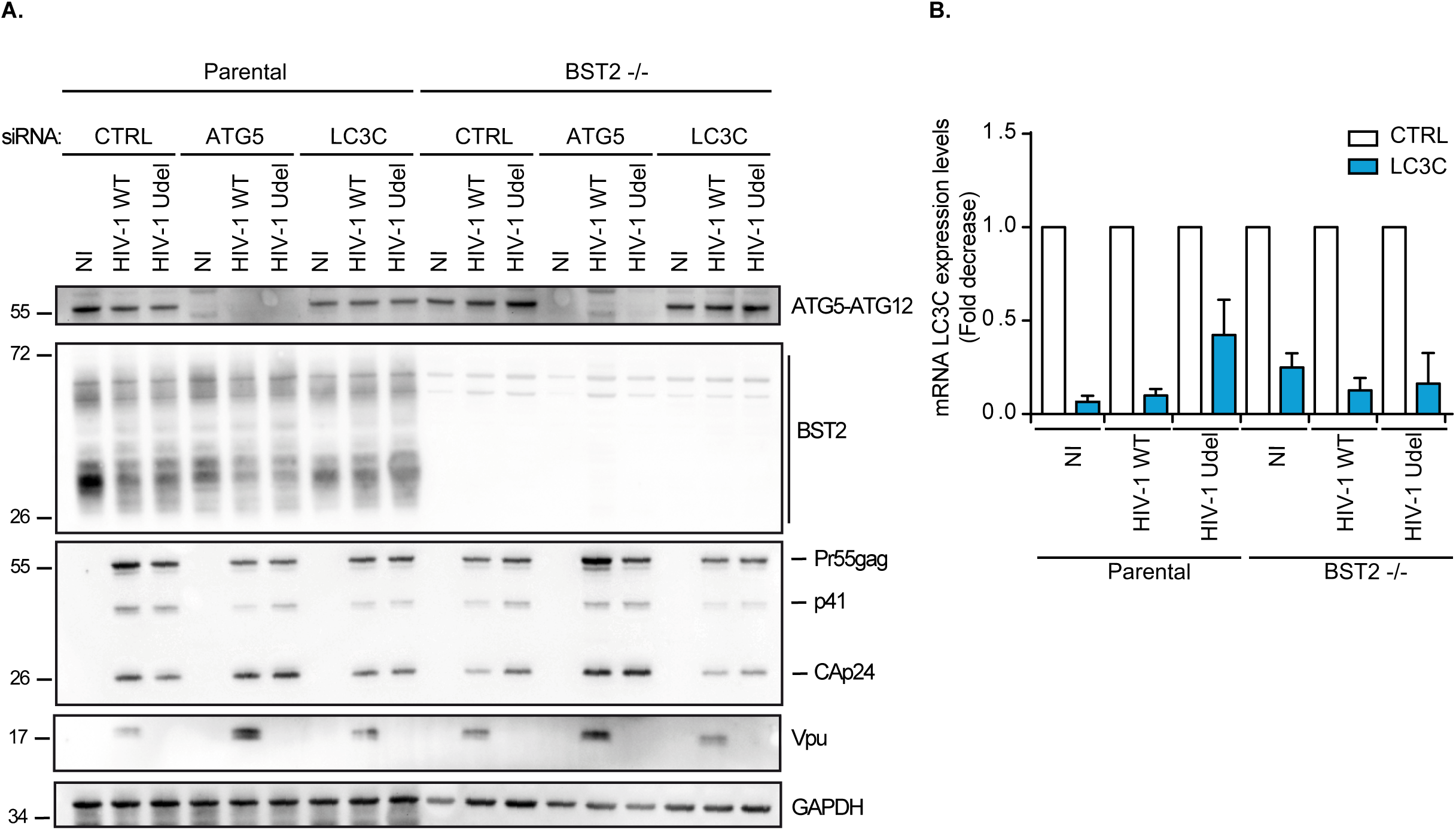
related to Figure 5. **(A)** Western blot analysis of ATG5, HIV-1 Gag and CAp24 products, BST2, Vpu and GAPDH in parental and BST2 -/- HeLa cells transfected with either control siRNA (CTRL) or siRNA targeting ATG5 (ATG5) or LC3C and transfected with the WT or Udel NL4.3 HIV-1 provirus for 28hrs. The western blot is representative of three independent experiments. **(B)** Total RNA profile for LC3C mRNA levels relative to KDSR by qRT-PCR of HeLa BST2 WT and HeLa BST2 -/- cells transfected with either control siRNA (CTRL) or siRNA targeting LC3C and transfected with the provirus HIV-1 NL4.3 WT or Udel for 28hrs. n= 3 experiments.

